# Reinforcement-Based Processes Actively Regulate Motor Exploration Along Redundant Solution Manifolds

**DOI:** 10.1101/2023.02.08.527668

**Authors:** Adam M. Roth, Jan A. Calalo, Rakshith Lokesh, Seth R. Sullivan, Stephen Grill, John J. Jeka, Katinka van der Kooij, Michael J. Carter, Joshua G. A. Cashaback

## Abstract

From a baby’s babbling to a songbird practicing a new tune, exploration is critical to motor learning. A hallmark of exploration is the emergence of random walk behaviour along solution manifolds, where successive motor actions are not independent but rather become serially dependent. Such exploratory random walk behaviour is ubiquitous across species, neural firing, gait patterns, and reaching behaviour. Past work has suggested that exploratory random walk behaviour arises from an accumulation of movement variability and a lack of error-based corrections. Here we test a fundamentally different idea—that reinforcement-based processes regulate random walk behaviour to promote continual motor exploration to maximize success. Across three human reaching experiments we manipulated the size of both the visually displayed target and an unseen reward zone, as well as the probability of reinforcement feedback. Our empirical and modelling results parsimoniously support the notion that exploratory random walk behaviour emerges by utilizing knowledge of movement variability to update intended reach aim towards recently reinforced motor actions. This mechanism leads to active and continuous exploration of the solution manifold, currently thought by prominent theories to arise passively. The ability to continually explore muscle, joint, and task redundant solution manifolds is beneficial while acting in uncertain environments, during motor development, or when recovering from a neurological disorder to discover and learn new motor actions.

## INTRODUCTION

When pushing a swinging door or grabbing a handrail, there are several potential locations we can place our hand. While such tasks appear simple, the sensorimotor system has the constant challenge of selecting an action from the infinite number of potential solutions along muscle,^1,2,3^ joint,^4,5,6,7^ and task redundant dimensions^8,9,10,11,12,13^ (e.g., grabbing a handrail). Past work has highlighted that humans are more variable along such redundant solution manifolds,^10,11,5,14,6,15,3,16^ which may reflect an exploratory mechanism that is continually searching for the most successful action.^17,18^ Continual exploration may be a beneficial strategy in a dynamic or uncertain environment.^19^

Promiscuous songbirds explore by injecting greater levels of variability in the pitch of their tune to attract several mates.^20,21,22,23,24^ Likewise, movement variability may facilitate the ability to find the most successful motor action.^17,18^ Movement variability has been proposed to arise from stochastic neuromuscular processes^25,26,27^ (‘motor movement variability’), the dorsal premotor cortex during movement preparation^8,28,29^ (‘planned movement variability’), and the basal ganglia^18,22,30^ (‘exploratory movement variability’). It has been proposed that the sensorimotor system has knowledge of both planned^31^ and exploratory^30,32^ movement variability, which arise from separate neural circuits. The role of the basal ganglia provides an explanation for the compromised regulation of exploratory movement variability for those with Parkinson’s Disease.^30^ It is unclear to what extent the sensorimotor system uses motor, planned, or exploratory movement variability to facilitate exploratory behaviour that promotes success.

Elegant theoretical and empirical work by van Beers and colleagues (2013) suggested that both knowledge of and acting upon planned movement variability leads to exploratory motor behaviour.^8,33,34,35^ They found that participants displayed greater explorative behaviour along the task-redundant dimension compared to the task-relevant dimension. Here the term exploration captures the ideas of increased variability as well as using their variability to traverse the solution manifold. They quantified exploration using statistical random walks (i.e., lag-1 autocorrelations), where a greater lag-1 autocorrelation is indicative of more exploration. In this context the term exploration captures not only the presence of movement variability, but also the idea that the sensorimotor system is aware of movement variability and allows it to update reach aim to traverse the solution manifold. Here greater exploratory random walk behaviour was attributed to passive process that arose from an accumulation of planned movement variability in the task-redundant dimension and not making trial-by-trial corrective actions based on error feedback. A fundamentally different explanation is that reinforcement-based processes may actively regulate the magnitude and structure of movement variability that underlies exploratory random walk behaviour—but this idea has not yet been tested empirically.

Here we hypothesize that reinforcement-based mechanisms contribute to exploratory random walk behaviour. To test this idea, we manipulated the size of both the visually displayed target and an unseen reward zone, as well as the probability of reinforcement feedback. For all three experiments we made *a priori* predictions with a general model. We then found the best-fit model from seven different plausible models, each representing a unique explanation of the mechanisms regulating sensorimotor exploration. Taken together, our empirical and modelling results support the idea that reinforcement-based mechanisms play a critical role in regulating exploratory random walk behaviour.

## RESULTS

### Experimental Design

In **Experiments 1** (n = 18), **2** (n = 18), and **3** (n = 18), participants made 500 reaching movements in the horizontal plane (**Fig. 1A**). For each trial, participants began their reach in a start position and attempted to stop within a virtually displayed target. They did not have vision of their hand. For each reach, we recorded their final hand position when they stopped within or outside the virtually displayed target.

**Figure 1:**
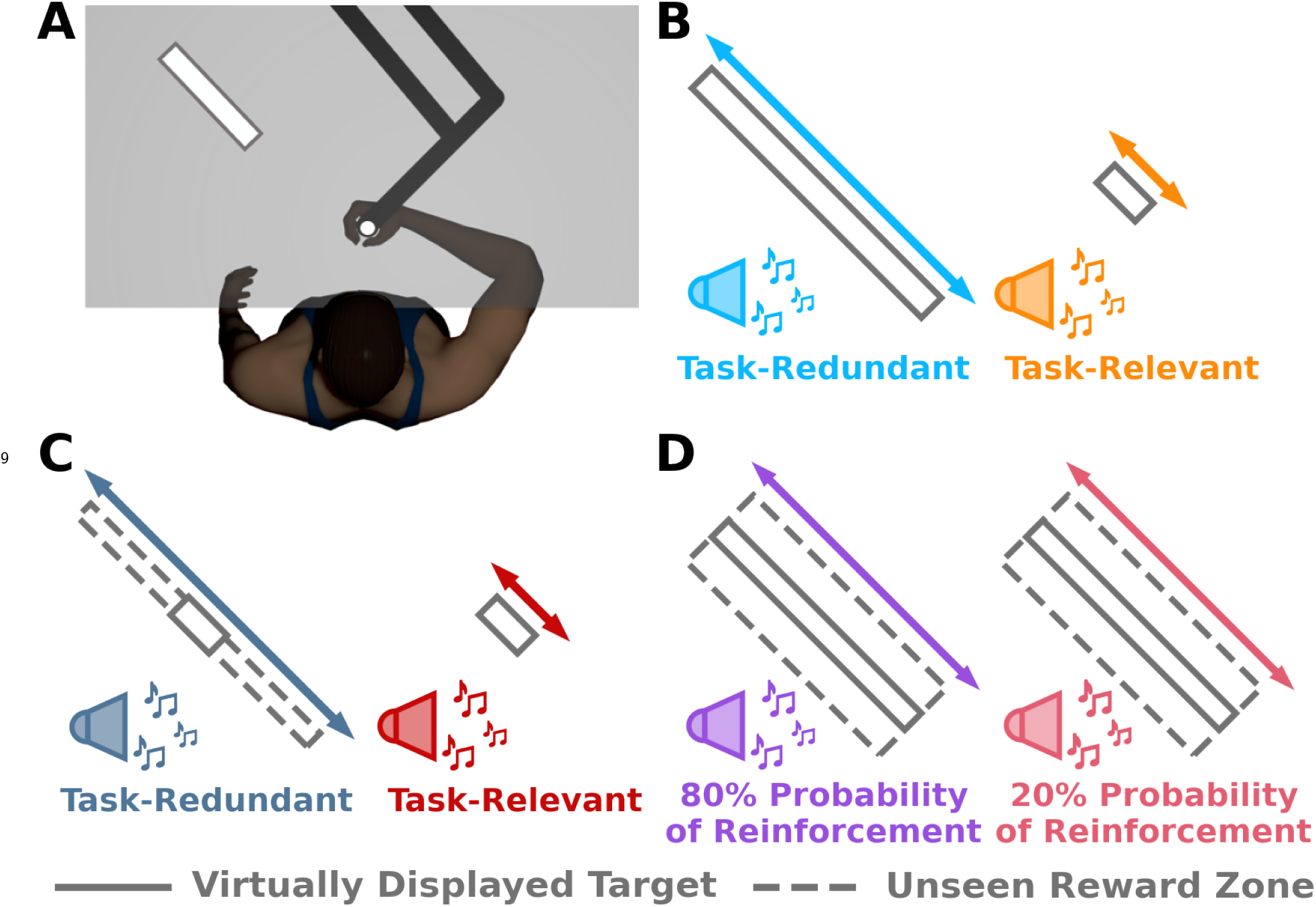
Apparatus and Experiment Design. **A)** Participants grasped the handle of a robotic manipu- landum. A semi-silvered mirror projected images (start position, target) from an LCD screen and occluded vision of the hand. Participants were instructed to reach from the start position (white circle) and attempt to stop their hand inside a virtually displayed target (white rectangle). For all experiments, participants were informed that they would receive positive reinforcement feedback (pleasant sound, target expands, small monetary gain; represented with the sound icons) if they successfully stopped within the target. **B) Experiment 1** examined the role of reinforcement feedback on exploratory sensorimotor behaviour. In separate blocks of trials, participants reached either to a long-rectangular target that promoted exploration along the movement extent (task-redundant, light blue) or a short-rectangular target (task-relevant, light orange) that discouraged exploration. Participants received positive reinforcement feedback if they stopped within the virtually displayed target. **C)** In **Experiment 2**, our goal was to control for the size of the virtually displayed target and to replicate the results of the first experiment. Participants always observed a short-rectangular target (solid grey outline) and received reinforcement feedback when their hand stopped within the reward zone (dashed grey outline). Critically, and unbeknownst to participants, in one condition there was a long-rectangular reward zone (task-redundant, dark blue) that promoted exploration. In the second condition, the reward zone matched the virtually displayed short rectangular target (task-relevant, dark red). **D)** In **Experiment 3** we manipulated the probability of reinforcement feedback while controlling for both the size of the virtually displayed target and the unseen reward zone. Participants were shown the same long rectangular target (dark grey outline) and probabilistically received positive reinforcement feedback when their hand stopped within the reward zone (grey dashed outline). There was an 80% probability (purple) and a 20% probability (pink) that participants would receive positive reinforcement feedback in either condition.

For all three experiments we used a repeated measures experimental design. Participants performed 50 baseline trials, 200 experimental trials, 50 washout trials, and then another 200 experimental trials. Condition order was counterbalanced for the experimental trials. During baseline and washout trials, participants reached towards and attempted to stop within a white circular target. For the first 40 trials of baseline and washout, a small cursor indicated final hand position. Participants received no feedback of their final hand position for the last 10 trials of baseline and washout. Removing feedback for the last 10 trials allowed us to estimate movement variability without the influence of reinforcement feedback or error feedback.

During the experimental trials, participants reached towards and attempted to stop within a rectangular target (**Fig. 1A**). The rectangular targets were positioned such that their major and minor axes corresponded to movement extent (i.e., parallel with the reaching movement) and lateral direction (i.e., orthogonal to the reaching movement), respectively. The target dimensions depended on the experimental condition and were scaled according to each participant’s movement variability from their last 10 baseline trials.^18^ Scaling target width based on baseline behaviour maintained a relatively constant task difficulty across participants. During the experimental trials, participants were informed that they would receive positive reinforcement when their final hand position was within the target, such that i) they would hear a pleasant sound, ii) the target would briefly expand, and iii) they would earn a small monetary reward. Participants were also informed they would not receive any feedback when they did not stop within the rectangular target.

In **Experiment 1**, we addressed how reinforcement feedback influences exploration along a redundant solution manifold. In the task-redundant condition, participants reached to a long rectangular target that promoted exploration along the movement extent (**Fig. 1B**). In the task-relevant condition, participants reached to a short rectangular target that discouraged exploration along the movement extent. Here we predicted greater explorative behaviour along the movement extent in the task-redundant condition compared to the task-relevant condition (see *A Priori Model Predictions* in the section below).

The goal of **Experiment 2** was to control for visual differences in the virtually displayed target size between conditions, since past work has shown that a visually larger target leads to greater movement variability.^36,13,37,38^ That is, we wanted to be assured that reinforcement-based processes were leading to greater exploration (lag-1 autocorrelation) rather than greater movement variability due to a larger visually displayed target. Further, this would also allow us to replicate the results from **Experiment 1**. Participants were shown the short rectangular (task-relevant) target in both conditions (**Fig. 1C**). They received reinforcement feedback when their hand stopped inside an unseen reward zone. In the task-relevant condition, the unseen reward zone matched the visually displayed target. Unbeknownst to participants, in the task-redundant condition they received reinforcement feedback if they stopped anywhere inside a long rectangular, unseen reward zone. Here we again predicted greater explorative bahaviour along the movement extent of the task-redundant target just as in **Experiment 1** (see *A Priori Model Predictions*).

In **Experiment 3** we directly manipulated reinforcement feedback to assess how it influences exploration along a redundant solution manifold. Specifically, we manipulated the probability of reinforcement feedback. By manipulating the probability of reinforcement feedback, we were able to simultaneously control for the size of both the visually displayed target and the unseen reward zone. Participants were always shown the task-redundant target. Unbeknownst to the participants, we widened the unseen reward zone in both conditions (**Fig. 1D**). Here, we manipulated the probability of receiving reinforcement feedback when participants stopped within the unseen reward zone. Participants performed two conditions, where they either had an 80% or 20% probability of receiving reinforcement feedback. We predict that participants will exhibit greater exploratory random walk behaviour with a higher probability of reinforcement feedback (see *A Priori Model Predictions*).

### A Priori Model Predictions

Only a few models of sensorimotor behaviour that simulate final hand position consider multiple sources of movement variability. Further, these models have varying assumptions on how much knowledge the sensorimotor system has of a particular source of movement variability when updating a motor action.^31,8,18,39,32^ Here we developed a general model (Model 1; **Eq. 1A,B**) that consolidated previously proposed models while using a minimal number of assumptions. We used our general model to generate *a priori* predictions. The general model simulates 2-dimensional final reach position (*X_t_*) and intended reach aim 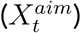 according to

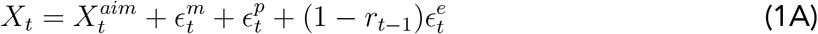

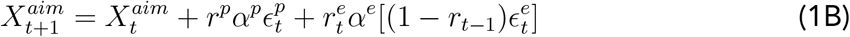

The model incorporates three sources of movement variability (*ϵ^i^*): motor (*ϵ^m^*), planned (*ϵ^p^*) and exploratory (*ϵ^e^*). Exploratory movement variability is added when the previous trial was unsuccessful^18,30,39^ 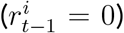. If the trial is successful 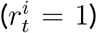, reach aim is updated proportionally (*α_p_,α_e_*) towards the planned and exploratory movement variability present in movement execution. Updating towards the previously reinforced motor actions results in a statistical random walk of the final hand positions. This random walk behavior is qualitatively described as exploration and is quantitatively captured using a lag-1 autocorrelation analysis. A greater lag-1 autocorrelation corresponds to greater exploratory random walk behavior. Note, while here we focus on sensorimotor exploration, this class of models can and have been used to capture sensorimotor adaptation.^31,8,18,39,32^

### Simulating Individual Behaviour

We first simulated the final hand position of an individual performing the task-redundant and task-relevant conditions from **Experiment 1** (**Fig. 2A**). Here we see more exploration, corresponding to a higher lag-1 autocorrelation, along the movement extent of the reach in the task-redundant condition (**Fig. 2B**). For this simulated individual, we then quantified the level of exploration along the movement extent for both conditions. A greater lag-1 autocorrelation indicates greater exploration of the solution manifold. Similar to past work,^8,9,34,12,40^ Model 1 produced greater lag-1 autocorrelation in the task-redundant condition compared to the task-relevant condition along the movement extent of the reach (**Fig. 2C**).

**Figure 2:**
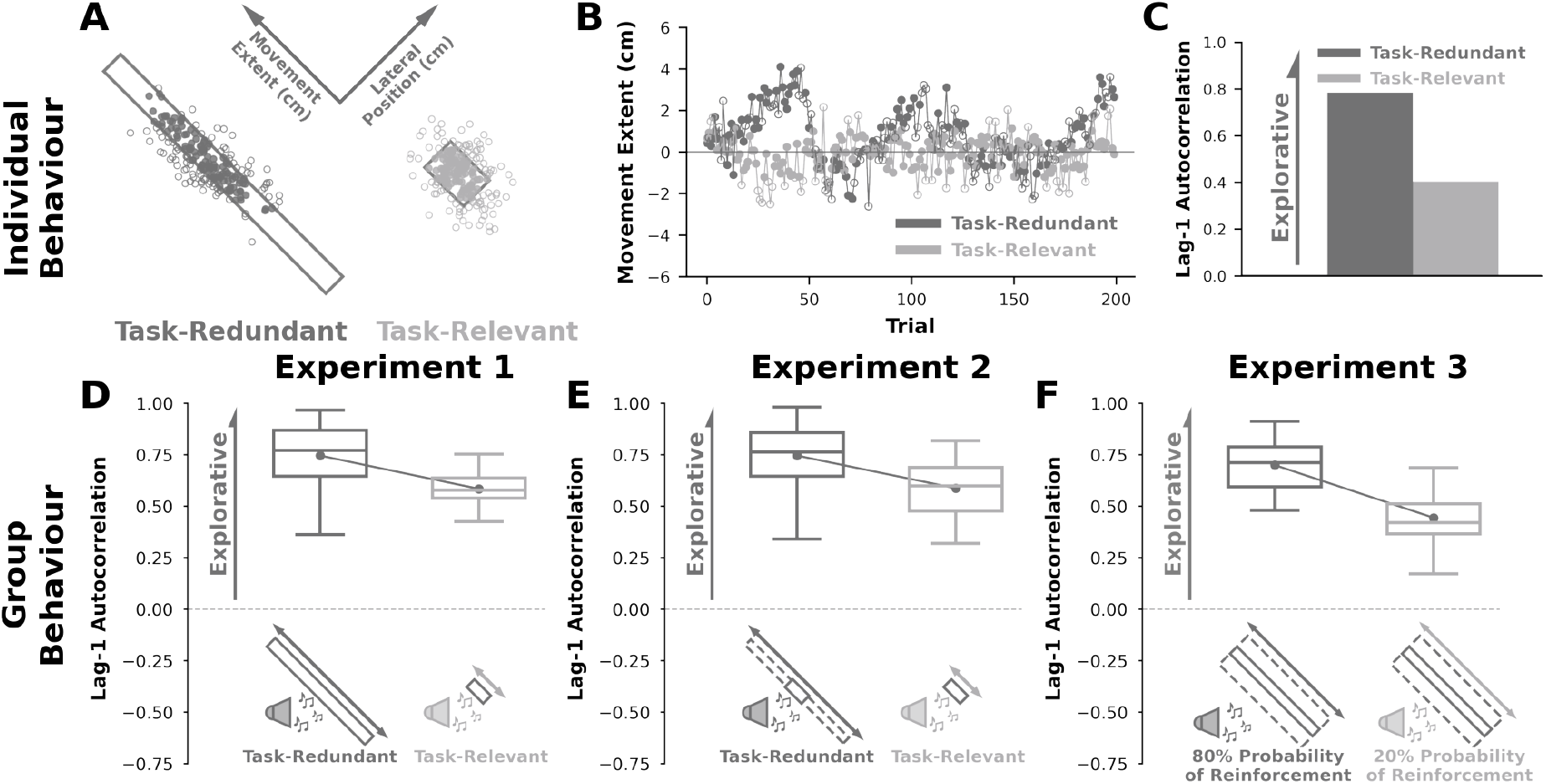
*A Priori* Model Predictions. We made theory-driven predictions by simulating (**A-C**) individual behaviour in **Experiment 1** and (**D-F**) group behaviour in **Experiments 1-3** using a model that updates reach aim based on reinforcement feedback while considering different plausible sources of movement variability (**Eq. 1**; Model 1). Model parameter values were held constant for all predictions. **A)** Successful (filled circle) and unsuccessful (unfilled circle) final hand positions when simulating an individual performing the task-redundant (dark grey) and task-relevant (light grey) conditions in **Experiment 1**. **B)** Corresponding final hand position (y-axis) for each trial (x-axis) along the major axes of the task-redundant and task-relevant targets. Note that there is greater exploration in the task-redundant condition. **C)** We quantified exploration by calculating the lag-1 autocorrelation (y-axis) of the trial-by-trial final hand positions along movement extent for each condition (x-axis). Here, a higher lag-1 autocorrelation represents greater exploration along a solution manifold. The model predicts greater lag-1 autocorrelation in the task-redundant condition (dark grey) compared to the task-relevant condition (light grey).**D-F)** Using the same parameter values, we simulated 18 participants per condition for the three experiments. **D)** For **Experiment 1**, the model predicted greater lag-1 autocorrelation (y-axis) in the task-redundant condition compared to the task-relevant condition (x-axis). **E)** In **Experiment 2**, the model predicted greater exploration in the task-redundant condition with a large rectangular unseen reward zone compared to the task-relevant condition. **F)** In **Experiment 3**, the model predicted greater exploration in the 80% probability of reinforcement feedback condition relative to the 20% probability of reinforcement feedback condition. Solid circles and connecting lines represent mean lag-1 autocorrelation for each condition. Box and whisker plots display the 25th, 50th, and 75th percentiles.

### Simulating Group Behaviour

To make *a priori* predictions of group behaviour for **Experiments 1**, **2**, and **3**, we used the general model (Model 1) to simulate the final hand position of 18 participants for each condition. For each condition, we calculated the trial-by-trial lag-1 autocorrelation for each participant. This analysis was performed separately along the movement extent and lateral direction of the reach. We used the average lag-1 autocorrelation for each condition as the *a priori* predictions for each experiment. *A priori* model predictions of group behaviour for **Experiment 1** (**Fig. 2D**) show higher lag-1 autocorrelations in the task-redundant condition compared to the task-relevant condition. Similarly, in **Experiment 2** the general model (Model 1) predicted greater lag-1 autocorrelation in the task-redundant condition in comparison to the task-relevant condition (**Fig. 2E**). In **Experiment 3**, the *a priori* model (Model 1) predicted greater lag-1 autocorrelation in the 80% probability of reinforcement condition compared to the 20% probability of reinforcement condition (**Fig. 2F**).

#### Experiment 1

In **Experiment 1**, we addressed how reinforcement feedback influences exploration along a task-redundant solution manifold. As a reminder, in the task redundant condition participants reached to a long rectangular target that was intended to promote exploration along its major axis (**Fig. 1B**). In the task-relevant condition, participants reached to a short rectangular target that discouraged exploration along its major axis.

**Fig. 3A** shows final hand positions for an individual participant in both the task-redundant and task-relevant conditions. This particular individual tended to reach towards the upper half of the task-redundant target, but the average final hand position across participants was within 1 cm of the target center. For this participant, we observed greater movement extent exploration of the solution manifold in the task-redundant condition (**Fig. 3B**). Conversely, we see less exploration in the task-relevant condition. Greater exploratory behaviour in the task redundant condition corresponded with a greater lag-1 autocorrelation (**Fig. 3C**).

**Figure 3:**
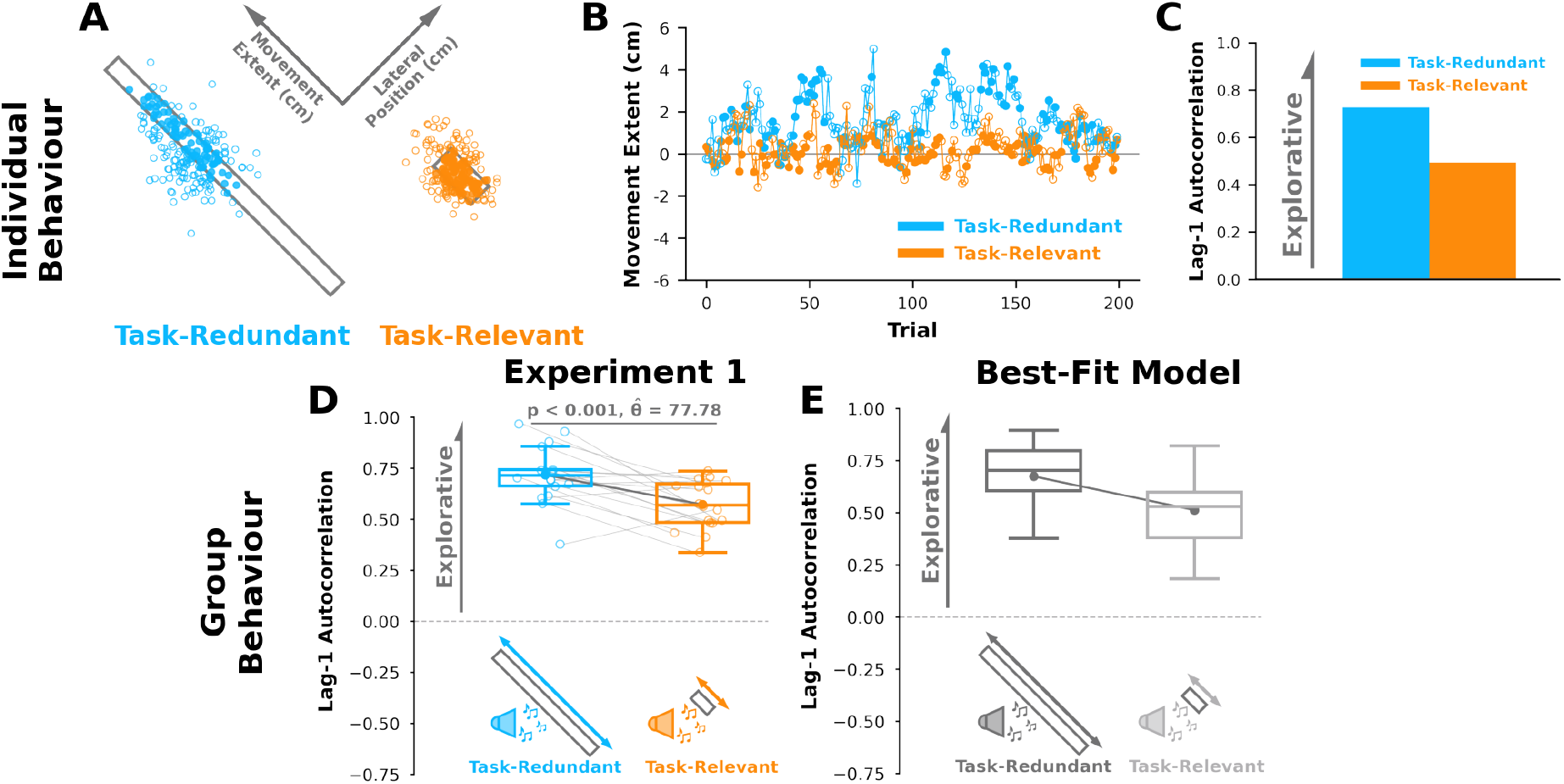
Experiment 1 Results. **A)** Successful (filled circle) and unsuccessful (unfilled circle) reaches by an individual participant performing the task-redundant (light blue) and task-relevant (light orange) conditions.**B)** Corresponding final hand position coordinates (y-axis) along the major axis of the task-redundant and task-relevant targets over trials (x-axis). **C)** This individual displayed greater lag-1 autocorrelation in the task-redundant condition, matching individual level model predictions (**Fig. 2C**). **D)** Participants (n = 18) had significantly greater lag-1 autocorrelation (p < 0.001) in the task-redundant condition (light blue) than the task-relevant condition (light orange), suggesting greater exploration of the task-redundant solution manifold. The group level data aligns with predictions of the *a priori* model (**Fig. 2D**). **E)** Best-fit model (**Eq. 4**; Model 4) according to both Bayesian Information Criteria (BIC) and Akaike Information Criteria (AIC) analyses (**Fig. 6A, B**). The best-fit model suggests that participants explored the task-redundant solution manifold by using exploratory movement variability and caching successful actions upon receiving reinforcement feedback. Solid circles and connecting lines represent mean lag-1 autocorrelation for each condition. Box plots represent the 25th, 50th, and 75th percentiles.

At the group level and aligned with our *a priori* model predictions (**Fig. 2D**), we see significantly greater lag-1 autocorrelation (p < 0.001, 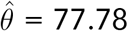) in the task-redundant condition compared to the task-relevant condition (**Fig. 3D**). These results suggest that participants explored the task-redundant dimension by updating their reach aim following positive reinforcement feedback.

Here we were primarily concerned with movement extent, which corresponded with our experimental manipulation along the major axis of the visually displayed target. We also examined lag-1 autocorrelations along the lateral direction that corresponded to the minor axis of the visually displayed targets. We did not see any lag-1 autocorrelation differences between conditions along the *lateral direction* (**Supplementary A, Fig. S1A**).

#### Experiment 2

Past work has shown that the distribution of final hand positions can be more variable when reaching to a large target compared to a small target,^36,13,37,38^ which could potentially influence exploratory behaviour. The goal of **Experiment 2** was to further test the idea that reinforcement feedback drove behaviour and replicate the findings in **Experiment 1**, while controlling for the visual size of the target. To control visually displayed target size, participants were only shown the short rectangular (task-relevant) target in both conditions. They received reinforcement feedback if they stopped within an unseen reward zone. In the task relevant condition, the unseen reward zone matched the visually displayed target. Critically, and unbeknownst to participants, in the task-redundant condition they received reinforcement feedback if they stopped anywhere inside a long rectangular, unseen reward zone (**Fig. 1C**).

Aligned with our *a priori* model predictions (**Fig. 2E**), participants displayed significantly greater lag-1 autocorrelation (p = 0.001, 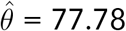) along the task-redundant condition compared to the task-relevant condition (**Fig. 4A**). These results replicate the finding that reinforcement feedback leads to greater exploratory behaviour along the task redundant dimension, while also highlighting that the results in **Experiment 1** were not due to visual size differences of the virtually displayed target.

**Figure 4:**
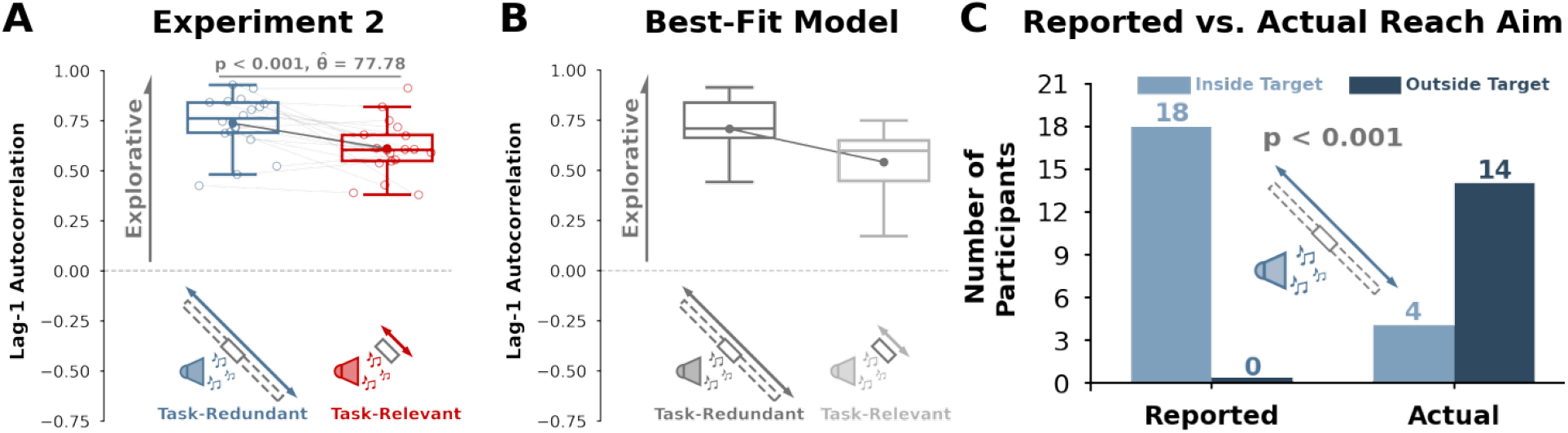
Experiment 2. **A)** Participants (n = 18) displayed significantly greater (p = 0.001) lag-1 autocorrelation (y-axis) in the hidden task-redundant condition (dark blue) compared to the task-relevant condition (dark red), despite observing the same visual target in each condition. The group behaviour aligns with *a priori* model predictions (**Fig. 2E**). **B)** Lag-1 autocorrelation (y-axis) for each condition (x-axis) based on simulations by the best-fit model (**Eq. 4**, Model 4). **C)** To test if they were aware of the long- rectangular reward zone in the task redundant condition (dark blue), participants were asked to mark on a target sheet where they were aiming for the task-redundant condition. All participants reported aiming to a point within the visual target (dark grey bars), but most participants had an average final hand position significantly outside the visual target (light green bars, p < 0.001). These data suggest that updating reach aim towards the most recent successful action may be in part driven by an implicit process (i.e., participants were unaware). Solid circles and connecting lines show mean lag-1 autocorrelation for each condition. Box and whisker plots show 25th, 50th, and 75th percentiles.

We assessed whether participants were aware of the reward zone manipulation in the task-redundant condition. After the experiment, participants were asked to mark their average final hand position for each condition on a sheet of paper that showed the task-relevant target. All participants reported that they were aiming somewhere within the visually displayed short rectangular target in the task-redundant condition. As a reminder, in the task-redundant condition participants saw the task-relevant target but received reinforcement feedback when they stopped within the task-redundant, unseen reward zone (**Fig. 1C**).

All 18 participants reported having an average final hand position within the visually displayed target. However, 14 participants had an average final hand position outside the visually displayed target (**Fig. 4C**, Fisher’s Exact Test, p < 0.001). These results suggest participants were unaware of the task-redundant, unseen reward zone and that updating reach aim may, in part, be driven by an implicit process.

#### Experiment 3

In **Experiment 3** we directly manipulated reinforcement feedback to further investigate its role in exploring task-redundant solution manifolds. Specifically, we manipulated the probability of reinforcement feedback. Critically, by manipulating only the probability of reinforcement feedback we simultaneously controlled for the size of both the visually displayed target and the unseen reward zone. Participants were always shown a task-redundant target (**Fig. 1D**). We manipulated the probability that the participants received reinforcement feedback if they stopped within the reward zone. Participants performed an 80% probability of reinforcement feedback condition and a 20% probability of reinforcement conditions.

Aligned with our *a priori* predictions (**Fig. 2F**), participants displayed significantly greater lag-1 autocorrelation (p = 0.002, 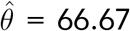) in the 80% probability condition compared to the 20% probability condition (**Fig. 5A**). This result suggests that partici-pants more frequently updated their reach aim when they received a higher probability of reinforcement feedback, which resulted in greater exploration of the solution manifold.

**Figure 5:**
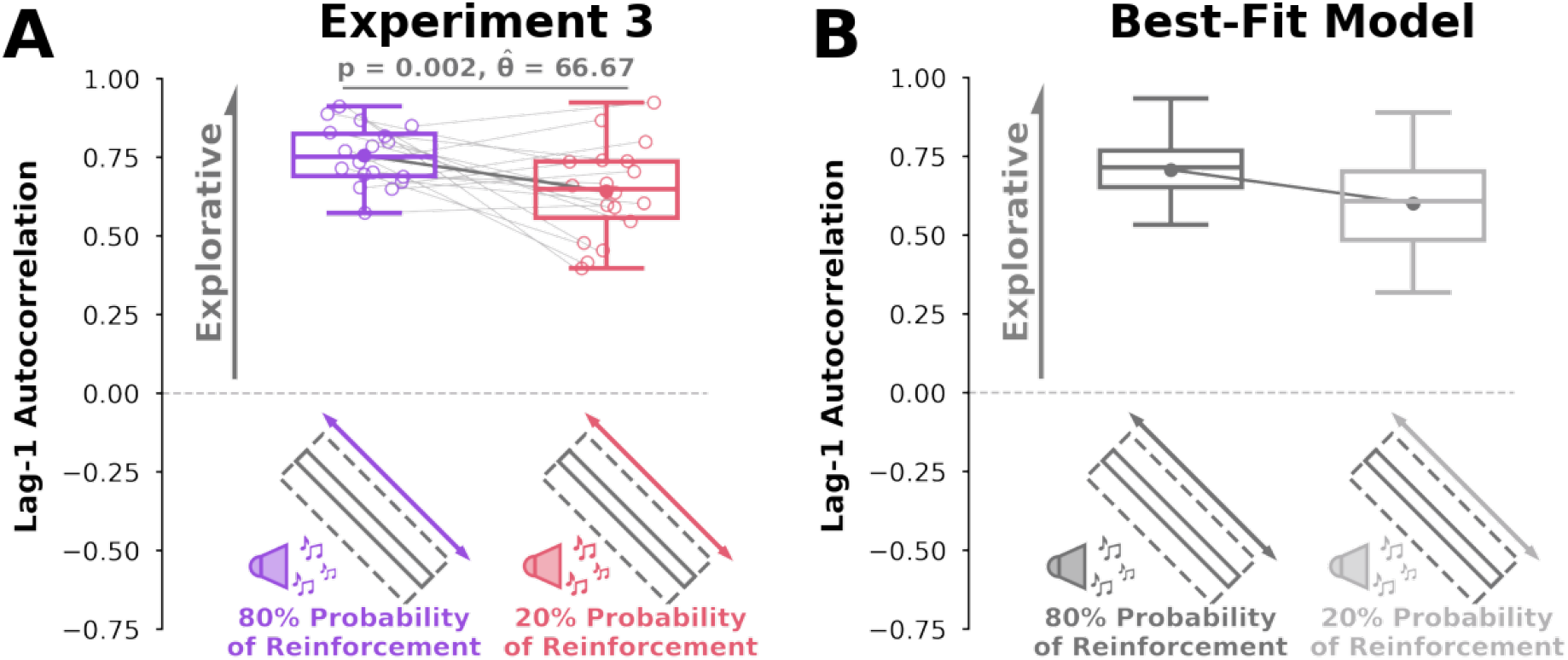
Experiment 3 Results. **A)** Participants (n = 18) displayed greater lag-1 autocorrelations (y-axis) with an 80% probability of receiving reinforcement feedback (purple) compared to a 20% probability of receiving reinforcement feedback (pink). Group behaviour aligned with *a priori* model predictions (**Fig. 2F**). **B)** Lag-1 autocorrelations (y-axis) for each condition (x-axis) based on simulations from the best-fit model. This model predicts greater lag-1 autocorrelation for the 80% probability of reinforcement feedback condition (purple) because there were more frequent reach aim updates with a higher probability of positive reinforcement feedback. Solid circles and connecting lines represent mean lag-1 autocorrelation for each condition. Box and whisker plots represent 25th, 50th, and 75th percentiles.

For each of our experiments, we wanted to control for the possibility that the observed lag-1 autocorrelations were the result of unknown stochastic processes rather than the sequential ordering of final hand positions based on reinforcement feedback. To test whether unknown stochastic processes caused the observed lag-1 autocorrelations, we performed a shuffling analysis^12,41^ (see **Supplementary B**). Our analysis suggests that the observed lag-1 autocorrelations are not the result of unknown stochastic processes (p > 0.05 for all participants).

### Best-Fit Model

Our general model (Model 1) consolidated previous reinforcement-based reaching models while considering multiple sources of movement variability. This general model did well to generate *a priori* theory-driven predictions. Yet there are other plausible mechanisms that the sensorimotor system uses to explore a redundant solution manifold. To test this idea, we considered four additional models (Models 2-5; **Eq. 2-5**, see **Methods**) by systematically reducing the number of free parameters from Model 1. Additionally, we tested two previously proposed reinforcement-based models (Models 6 and 7, see **Methods**) in the literature.^18,32^

Each model was simultaneously fit to each experimental condition across the three experiments (see **Methods**). We used Bayesian Information Criteria (BIC) and Akaike Information Criteria (AIC) analyses to compare the fit of each model while penalizing additional free parameters. For both BIC and AIC analyses, a lower score indicates a more plausible model. In agreement, both the BIC (**Fig. 6A**) and AIC (**Fig. 6B**) analyses supported Model 4 (**Eq. 4A,B**) as the Best-Fit Model. The best-fit model (Model 4) simulates final reach position and intended reach aim as follows

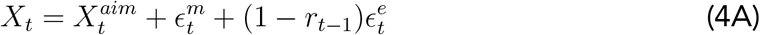

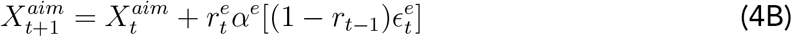

**Figure 6:**
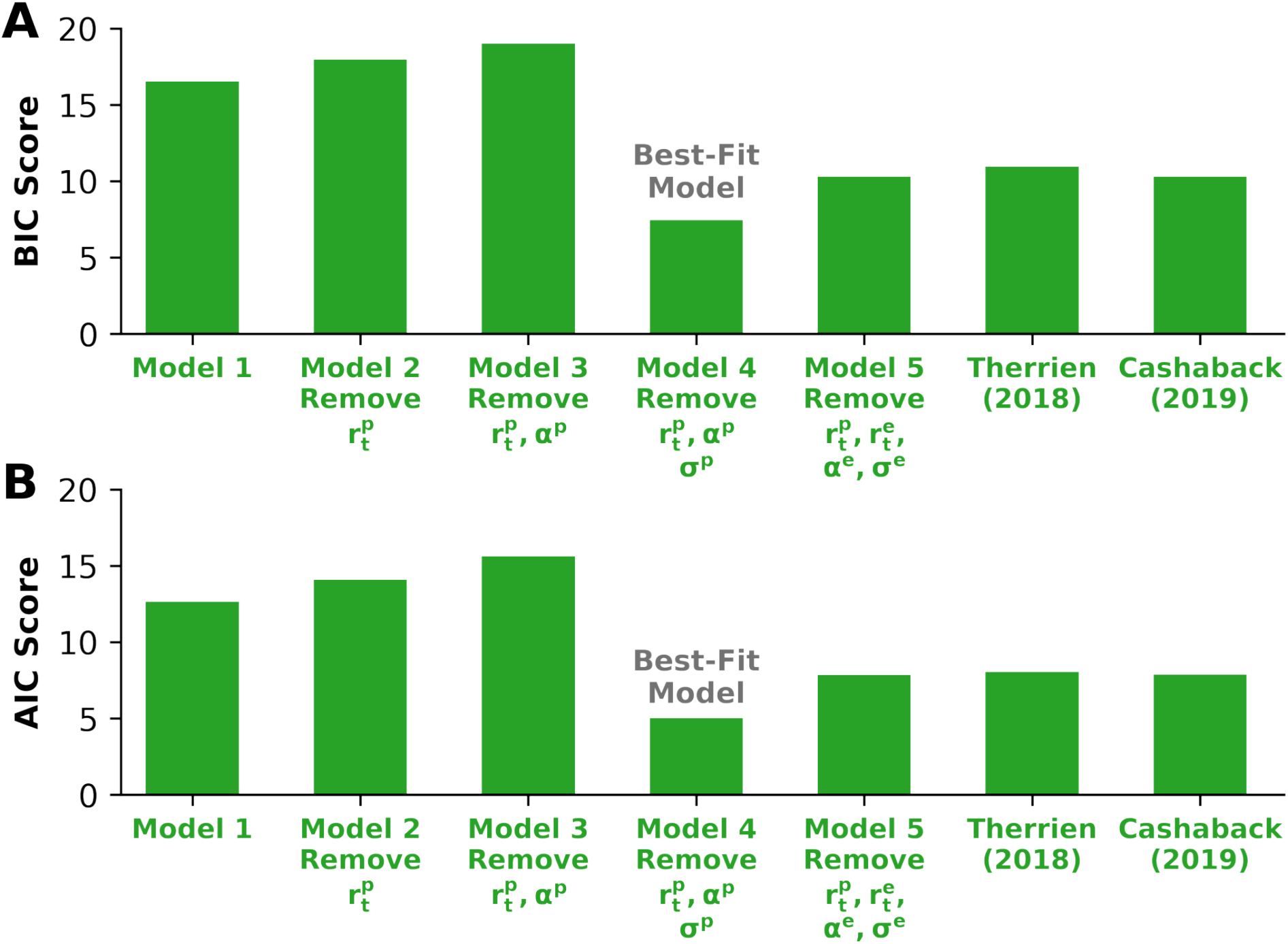
Model Selection. To compare each possible model (x-axis), we used **A)** Bayesian Information Criteria (BIC) and **B)** Akaike Information Criteria (AIC) analyses. For both analyses, a lower score represents a better model that strikes a balance between increasing the quality of the fit and decreasing the number of free parameters. Both analyses support removing all planning terms 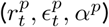 as the best-fit model. The best-fit model expands its movement variability after unsuccessful reaches in an attempt to find a successful reach position. The model updates its intended reach aim towards the previously successful reach. This process of expanding movement variability following failure and updating towards successful reaches results in continual exploration of the task solution manifold.

Unlike the general model, the best-fit model (Model 4) does not consider planned movement variability. The model updates its aim towards the next successful final hand position 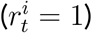 following an unsuccessful trial 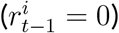. This suggests that the sensorimotor system uses knowledge of exploratory movement variability to update reach aim when behaviour is driven by reinforcement feedback.

Using our bootstrapping procedure, we obtained posterior distribution estimates of the model parameters (**Supplementary C**). We used the median values of the parameter posterior distributions to simulate participant reaching behaviour for **Experiment 1** (**Fig 3E**), **Experiment 2** (**Fig. 4B**), and **Experiment 3** (**Fig. 5B**). The model did well to capture lag-1 autocorrelations across all three experiments along the movement extent and lateral direction (**Supplementary A; Fig. S1**).

### Movement Variability is Greater with Failure

Past work has shown that binary reinforcement feedback modulates movement variability.^18,42,30,43,44,39,45,46^ An assumption of our models is that an unsuccessful reach leads to an increased amount of exploratory movement variability. To test this assumption, for each condition we calculated movement variability independently for trials following reinforced and unreinforced reaches. We observed greater movement variability following unreinforced trials compared to positively reinforced trials across all three experiments (p < 0.001 for all experimental conditions; **Supplementary D, Fig. S4**).

Recent literature has examined the effects of reinforcement (reward) feedback and task success on movement vigor.^47,48,49,50,51^ We examined reaction times as a proxy of movement vigor (see **Supplementary E**). However, we did not find consistent evidence to suggest that reinforcement feedback modulated reaction times in our experiments.

### The Magnitude and Structure of Movement Variability are Experimentally Dissociable and Predicted by the Best-Fit Model

Past studies have quantified exploration by comparing the relative variability between task-redundant and task-relevant dimensions^4,11,5,14,6,7^ (i.e., ‘uncontrolled manifold’ and ‘orthogonal dimension’). For each condition, we quantified the magnitude of movement variability by calculating the interquartile range (IQR) of participant final hand positions along the movement extent. In **Experiment 1** (**Fig. 7A**) and **Experiment 2** (**Fig. 7B**), we took the IQR ratio between the task-redundant and task-relevant for each participant. Participants displayed an IQR ratio significantly greater than one in **Experiment 1** (p = 0.034) and **Experiment 2** (p = 0.047), such that there was a greater magnitude of movement variability in the task-redundant conditions relative to the task-relevant conditions. In **Experiment 3** (**Fig. 7C**), we took the IQR ratio between the 80% and 20% probability of reinforcement conditions. Participants did not display an IQR ratio greater than one in **Experiment 3** (p = 0.296), where the magnitude of movement variability was similar between conditions.

**Figure 7:**
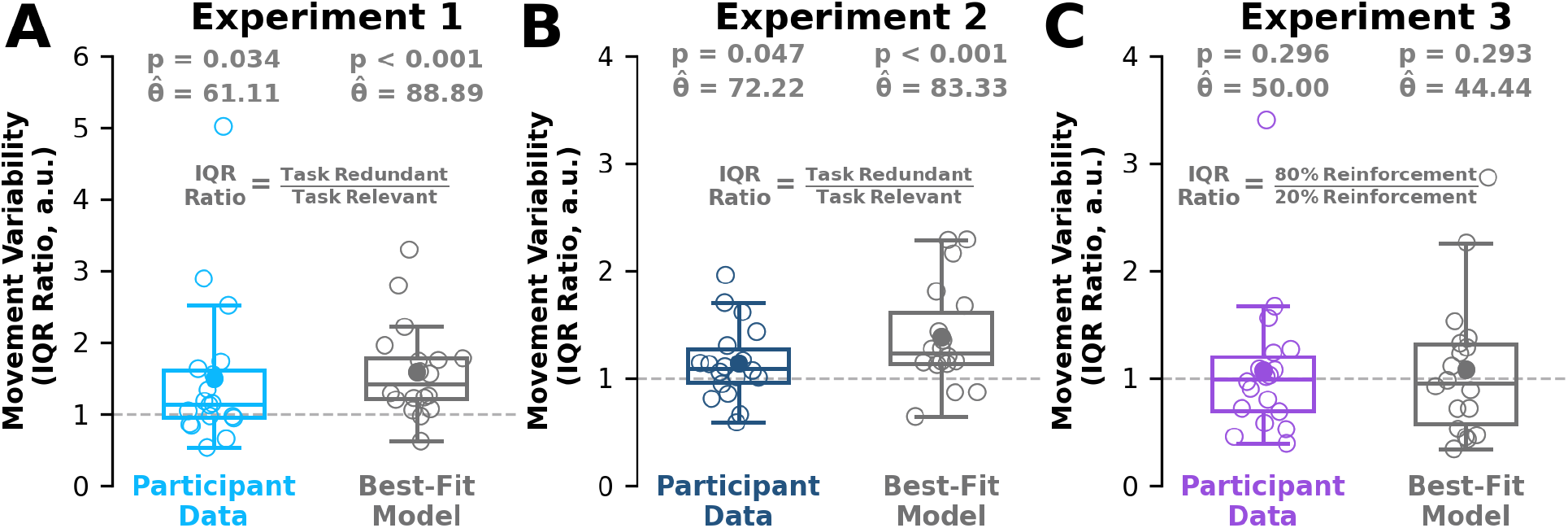
Interquartile Range Ratio. **A,B,C)** We calculated the interquartile range (IQR) of final hand positions along movement extent for each condition. Here we show the IQR ratio between conditions (y-axis) for participant data (colored) and best-fit model simulations (dark grey) for each experiment. Hollow circles represent individual data. Solid circles represent the mean IQR ratio. Box and whisker plots represent 25th, 50th, and 75th percentiles. An IQR ratio greater than one (dashed grey line) indicates greater movement variability (**A,B**) along the task-redundant condition or (**C**) the 80% probability of reinforcement condition. Participants displayed an IQR ratio greater than one in **A) Experiment 1** (light blue) and **B) Experiment 2** (dark blue). Thus, they had greater variability in their final hand positions during the task-redundant conditions compared to the task-relevant conditions. Participants in **C) Experiment 3** (purple) did not display an IQR ratio significantly greater than one. This suggests that participants showed similar levels of movement variability between the 80% and 20% probability of reinforcement conditions. Our best-fit model (Model 4, dark grey) replicated the observed data in **A) Experiment 1**, **B) Experiment 2**, and **C) Experiment 3**. Our results suggest that the magnitude (IQR ratio) and structure (lag-1 autocorrelation) of movement variability are dissociable between conditions and should be considered in tandem when assessing exploratory sensorimotor behaviour.

Interestingly, our analysis on the magnitude of movement variability is dissociated from the structure of movement variability (i.e., lag-1 autocorrelation analysis), where in the latter we see differences between conditions in all three experiments. This suggests a dissociation between the magnitude (IQR) and trial-by-trial structure (random walk) of movement variability between conditions. We performed the same IQR movement variability analysis using our best-fit model (Model 4) to test whether this dissociation can be explained by updating reach aim using exploratory movement variability after a successful trial. The best-fit model captures the observed trends in **Experiment 1** (**Fig. 7A**, p < 0.001), **Experiment 2** (**Fig. 7B**, p < 0.001), and **Experiment 3** (**Fig. 7C**, p = 0.320). This finding was consistent when we used the best-fit model to simulate 1,000,000 trials per condition.

Taken together, our behavioural and modelling results support the idea that reinforcement-based mechanisms play an important role in driving exploratory behaviour along task-redundant solution manifolds.

## DISCUSSION

We show that reinforcement feedback regulates sensorimotor exploration along task-redundant solution manifolds. Our finding was robust across a series of experiments where we manipulated the size of both the visually displayed target or unseen reward zone, as well as the probability of reinforcement. Our work suggests that exploratory random walk behaviour arises from utilizing knowledge of movement variability to update intended reach aim when an action is positively reinforced—leading to active and continual exploration of the solution manifold. This mechanism can also explain an observed experimental dissociation between the magnitude and structure of movement variability.

In this paper we aimed to address whether reinforcement feedback contributed to exploratory random walk behaviour. In **Experiment 1**, we found that participants displayed greater exploratory random walk behaviour along the task-redundant dimension compared to a task-relevant dimension when receiving positive reinforcement feedback that indicated success. Others have reported that a visually larger target leads to greater movement variability.^36,13,37,38^ Thus, to be assured a visually larger target alone was not causing changes in exploratory random walk behaviour we collected a second experiment. In **Experiment 2** we held the visually displayed target size constant and individuals received reinforcement when their final hand position was within a task-relevant or task-redundant reward zone. Replicating the results of **Experiment 1**, we again found greater exploration in the task-redundant dimension. For the task-redundant condition, we also found that participants were unaware of the large, unseen reward zone relative to the smaller visually displayed target, which may suggest an implicit role of reinforcement-based processes during exploratory behaviour. In **Experiment 3**, we directly manipulated reinforcement feedback while holding both the visually displayed target and unseen reward zone dimensions constant. Participants exhibited a greater trial-by-trial, exploratory random walk behaviour with a higher probability of reinforcement feedback. Collectively, these findings support the idea that reinforcement feedback plays an important role in regulating exploratory random walk behaviour.

In a previous study by van Beers and colleagues (2013), participants reached to a long rectangular target and received vectored error-based feedback of their final hand position via a cursor. They also received a numerical reward score that scaled as a function of their distance from the target. They found that participants displayed exploratory behaviour (lag-1 autocorrelation ≈ 0.55) along the task-redundant dimension of a rectangular target. Conversely, along the task-relevant dimension of the rectangular target they had significantly less exploration (lag-1 autocorrelation ≈ 0.0), aligning with making corrective actions based on error-based feedback.^31,33^ The authors attributed greater explorative behaviour along the task redundant dimension to passively allow an accumulation of planned movement variability and a lack of error-based corrections.

In their model, behaviour was explained by adjusting the error correction term separately for the task redundant and task-relevant dimensions. An alternate, yet potentially complementary idea, is that exploration along task-redundant solution manifolds are driven by reinforcement-based processes. We recently proposed theoretical work that suggests reinforcement-based processes could also explain exploratory random walk behaviour,^18^ but this idea had not yet been tested empirically. Aligned with this idea, across our three experiments we show that reinforcement-based processes contribute to exploratory random walk behaviour along task-redundant dimensions. Interestingly, we observed greater exploration along task-redundant dimensions when behaviour was driven by reinforcement feedback relative to past work that used error feedback.^8^ Similar to past adaptation studies,^52^ an interesting future direction would be to examine the individual roles and interplay between reinforcement-based and error-based processes during sensorimotor exploration.

It has been proposed that the sensorimotor system has knowledge of both planned movement variability that arises in the dorsal premotor cortex^8,28,29^ and exploratory movement variability that arises in the basal ganglia.^20,53,22^ An accumulation of either of these processes could lead to exploratory random walk behaviour. Critically, however, planned movement variability that is not conditioned on positive reinforcement could not explain the observed differences in **Experiment 1, 2**, or **3**. For example, Model 5 accumulates planned movement variability every trial regardless of reinforcement feedback and was unable to capture differences between conditions. Conversely, models (i.e., Model 1, Model 3, Model 4, Cashaback 2019, Therrien 2018) that included knowledge of exploratory movement variability to update reach aim conditioned on reinforcement feedback could capture the observed trends. Similar to others, we found greater exploratory movement variability following failure (**Supplementary D, Fig. S4**). Exploratory movement variability has been closely linked to the basal ganglia and the dopaminergic system,^22^ which scales with reward prediction error in rodents.^54^ In humans, those with Parkinson’s disease become unable to regulate movement variability as a function of reinforcement feedback.^30^ Parkinson’s disease may be a population of interest to gain causal insight into the influence of reward prediction error and other reinforcement-based processes on sensorimotor exploration.

One of the seven plausible models we considered aligned with the use-dependency hypothesis (Model 5). The use-dependency hypothesis suggests that the sensorimotor system biases a movement to be similar to the previous movement.^55,56^ The use-dependency hypothesis also suggests that this behavioural change is not conditioned on reinforcement-based processes. The model proposed by van Beers^8^ in some respects resembles the use-dependency hypothesis along the task redundant dimension. In their model planned movement variability accumulates along the task-redundant dimension, which biases a movement to be similar to the previous movement. Similarly, our model 5 used planned movement variability, not exploratory movement variability, but failed to capture lag-1 autocorrelation differences between conditions in **Experiment 1, 2**, or **3**. These model results imply that the differences in exploratory behaviour between conditions is not driven by a use-dependency mechanism. A more parsimonious account of our findings, captured in our general model (Model 1) and the best-fit model (Model 4), is that the sensorimotor system will bias a movement when the previous movement is conditioned on reinforcement feedback. Further, it calls into question whether planned and exploratory movement variability arising in the dorsal premotor cortex and the basal ganglia are unrelated processes. Beyond performing model comparisons it is difficult to separate the relative contributions of use-dependency and reinforcement-based mechanisms in the current experiment paradigm. For example, use-dependency may to some extent contribute to baseline levels of exploratory statistical random walk behaviour. It would be useful to test whether use-dependency is conditioned on intrinsic or extrinsic reinforcement-based processes. Nevertheless, our results suggest that a reinforcement-based mechanism is necessary to explain the trends observed across all three experiments.

After **Experiment 2**, we asked participants to indicate their average final hand position for each condition. By asking after each condition, we avoided impacting exploration during the experiment through the use of self-reports of trial-by-trial hand positions that have been shown to alter behaviour.^57,58,59,60^ Our approach is admittedly a weak assessment of implicit processes that may contribute to exploratory behaviour. However, our results align with previous work suggesting participants are unable to localize their hand position when receiving only binary reinforcement feedback.^61^ Holland and colleagues (2018) showed that participants can develop and remove explicit strategies when receiving binary reinforcement feedback in a visuomotor rotation task. However, when participants were asked to remove their explicit strategy during washout, their reach angles did not fully return to baseline levels. Slightly elevated reach angles during washout may to some extent reflect an implicit component of reinforcement-based processes. It would be useful to examine whether there is some level of implicit reinforcement-based processes that contribute to exploratory sensorimotor behaviour.

An observation in the literature is that there is more movement variability along muscle,^1,2,3^ joint,^4,5,6,7^ and task^8,9,10,11,12,13^ solution manifolds. This observation has been classically studied in joint space using a dimensionality reduction technique proposed by proponents of the uncontrolled manifold hypothesis^7^ (UCM). The UCM hypothesis posits that humans have increased variability along redundant dimensions that have little impact on task success. This observation can be explained in the context of optimal feedback control, where biological systems tend not to intervene along redundant dimensions because it is energetically costly (termed the ‘minimum intervention principle’).^15^ In other words, the observed increases in movement variability have been previously assumed to occur passively by not intervening. Alternatively, our results show that an increase in variability along task-redundant dimensions may occur from reinforcement-based processes that actively and continually update reach aim towards recently successful motor actions. Here we analyzed both the magnitude (interquartile range) and trial-by-trial structure (lag-1 autocorrelation) of the movement variability along the movement extent of the reach between conditions. Across **Experiments 1**, **2**, and **3**, we see that the trial-by-trial structure of movement variability (lag-1 autocorrelation) changes depending on unseen reward zone size or the probability of reinforcement feedback. However, the magnitude of movement variability (interquartile range) only changes with the unseen reward zone size and not the probability of reinforcement feedback. These results suggest a dissociation between the magnitude and structure of movement variability between conditions. While the magnitude of movement variability analysis aligns with both the UCM hypothesis and OFC, neither framework captures changes in the trial-by-trial structure of movement variability. In contrast to past theories, our work suggests that reinforcement-based processes lead to the active and continual exploration of task-redundant solution manifolds.

Past work, including our own, has considered increases in trial-by-trial movement variability following failure as a metric of exploration^39,62,18,30^ (**Supplementary D**). Likewise, we also considered the distribution of final hand positions (i.e. interquartile range) across several trials to assess movement variability. Both trial-by-trial and interquartile range assessments of movement variability only describe a single aspect of the exploratory process. Greater movement variability does not necessarily imply exploration or lead to improved performance. Critically, participants must also have knowledge and act upon movement variability to update their reach aim to increase the likelihood of producing a successful motor action. Lag-1 autocorrelation provides a metric to assess whether the sensorimotor system has knowledge of and acts upon movement variability to explore. Collectively our results show that the sensorimotor system modulates movement variability as a function of reinforcement feedback, has knowledge of movement variability, and acts upon movement variability to update reach aim following positive reinforcement. Thus, exploration can be considered as a feedback modulated process of expanding and utilizing knowledge of movement variability to actively and continually explore the solution manifold.

Exploratory random walk behaviour has been universally seen across species,^8,34,22^ along neural manifolds,^34,63^ gait cycles,^12,41^ and trial-by-trial reaching behaviour.^8,33,9,40^ Humans show greater movement variability along muscle,^1,2,3^ joint,^4,5,6,7^ and task^8,9,10,11,12^ solution manifolds, which to some extent may be driven by reinforcement-based processes continually exploring for the best possible action. Across three experiments, we showed evidence to suggest that exploratory random walk behaviour arises from utilizing knowledge of exploratory movement variability to update intended reach aim when an action is positively reinforced. This mechanism leads to active and continual exploration that is useful for finding successful motor actions in dynamic or uncertain environments. The ability to explore is also particularly relevant following a musculoskeletal or neurological disorder, where a new set of actions must be discovered and learned to perform everyday, functional tasks.

## METHODS

### Participants and Power Analyses

We collected 54 participants across **Experiment 1** (n = 18, 20.4 years ± 2.7 SD), **Experiment 2** (n = 18, 20.6 years ± 2.2 SD), and **Experiment 3** (n = 18, 20.8 years ± 2.9 SD). We used a power analysis, with a smallest effect size of interest of *d* = 0.5 (repeated measures), *α* = 0.05 (one-tailed based on theory-driven predictions), and *β* = 0.2 (80% statistical power) to find the number of participants. The power analysis suggested that 17 participants were needed for sufficient statistical power. However, we recruited 18 participants per experiment to properly counterbalance each experiment. Participants reported they were right-handed and free of neuromuscular disease. Each participant provided informed consent to procedures approved by the University of Delaware’s In ternational Review Board.

### Apparatus

Participants grasped the handle of a robotic manipulandum (**Fig. 1A**, KINARM, BKIN Technologies, Kingston, ON, Canada) and made reaching movements in the horizontal plane. A semi-silvered mirror blocked vision of both the participant’s upper-limb and the robotic arm, and projected images (start position, target) from an LCD screen onto the horizontal plane of motion. Hand position was recorded at 1000 Hz and stored offline for data analysis.

### General Task Protocol

Participants were presented with virtual images of a start position (white circle, radius = 0.75cm) that was aligned with the sagittal plane and approximately 15 cm away from their body. Displayed targets were located 45 degrees to the left of the sagittal plane and 15 cm away from the start position (**Fig. 1A**). Rectangular targets were rotated so that their major axis aligned with the center of the start position. For each trial, participants began from the start position and were instructed to “reach and stop inside the target.” The start position turned yellow after a short, randomized delay (250-1000 ms) to signal the beginning of the trial. Final hand position was defined as the participant’s hand location 100 ms after their hand velocity went below 0.045 cm/s. One second after stopping, the robot used a minimum jerk trajectory to return their hand to the start position.

For all three experiments, participants performed 50 baseline trials, 200 experimental trials, 50 washout trials, and then another 200 experimental trials. During baseline and washout trials, participants reached towards and attempted to stop within a white circular target (radius = 0.5 cm). Participants saw a small cursor (radius = 0.25 cm) appear at their final hand position for the first 40 baseline and 40 washout trials. No feedback was given for the final 10 baseline and final 10 washout trials. During the experimental trials of the first condition (trials 51-250) and second condition (trials 301-500), participants reached towards rectangular targets. We told participants that they would receive the following positive reinforcement feedback each time they stopped within the target: 1) hear a pleasant sound, 2) the target would briefly expand, and 3) earn a small monetary reward.

Participants were told that base compensation was $5.00 USD and they could earn an additional $5.00 USD performance bonus based on task performance. After completing the experiment, all participants received $10.00 USD irrespective of task performance.

### Experiment 1 Design

In **Experiment 1**, we addressed how reinforcement feedback influences exploration of task-redundant solution manifolds. Past work^8^ addressing exploration of redundant dimensions in a reaching task compared exploration along movement extent (i.e., parallel with the reaching movement) and lateral direction (i.e., orthogonal to the reaching movement). Movement variability can vary between the movement extent and lateral direction,^64,65^ which may impact exploratory behaviour. To control for potential differences between the two dimensions, we designed the task such that all comparisons are made along the movement extent of a reach.

In the first condition, participants reached to a long rectangular (task-redundant) target (**Fig. 1B**). The major axis length of the task redundant target was 12 cm.^8^ During the second condition, participants reached to a short rectangular (task-relevant) target. For each participant, the major axis length of the task-relevant target was proportional to their movement variability along the extent of the reach during the last 10 baseline trials (1.4σ, 2.1 ± 0.14 cm). We expected to find greater exploration along the task-redundant target compared to the task-relevant target. The minor axis length of the task-redundant (0.95σ, 1.0 ± 0.09 cm) and task-relevant (1.4σ, 1.6 ± 0.13 cm) was proportional to each participant’s lateral movement variability during the last 10 baseline trials. The slight differences in minor axis length between the two conditions (5.6 mm) was designed to promote similar levels of success, which has been shown to influence motivation.^66^ Condition order was counterbalanced.

### Experiment 2 Design

The goal of **Experiment 2** was to control for visual target size between conditions while replicating the findings of **Experiment 1**. Participants performed two conditions. In both conditions, participants were only shown the task-relevant target (**Fig. 1C**). The visually displayed task-relevant target was proportional to each participant’s movement variability during the last 10 baseline trials, as described in **Experiment 1**. Unbeknownst to participants, in one condition they received positive reinforcement feedback if they stopped anywhere inside an unseen reward zone. This reward zone had the same dimensions as the task-redundant target used in **Experiment 1**. In the second condition, the unseen reward zone matched the task relevant, visually displayed target. We expected to find greater exploration in the condition that used the task-redundant, unseen reward zone. Condition order was counterbalanced. After the experiment, participants were asked to mark their average reach position for each condition on a sheet of paper that showed the task-relevant target.

### Experiment 3 Design

In **Experiment 3** we investigated how the probability of reinforcement influences exploration of redundant task spaces, while controlling for the size of both the visually displayed target and the unseen reward zone. Participants were shown the task-redundant target. The dimensions of the visually displayed task-redundant target were the same as described in **Experiment 1**. Unknown to participants, we expanded the reward zone width in both conditions (**Fig. 1D**). For each participant, the unseen reward zone width was proportional to their lateral movement variability during the last 10 baseline trials (20, 2.59 ± 0.24 cm). Critically, we manipulated the probability of receiving positive reinforcement feedback when a participant stopped inside the unseen reward zone. In the first condition, participants had an 80% probability of receiving positive reinforcement feedback. In the second condition participants had a 20% probability of receiving reinforcement feedback. We expected to find greater exploration in the 80% condition compared to the 20% condition. Condition order was counterbalanced.

### Data Analysis

We performed data analysis using custom Python 3.8.12 scripts. For all experiments, final hand position coordinates were projected onto a rotated cartesian coordinate system that was aligned with the major and minor axes of the rectangular targets. Thus the x-axis and y-axis of the rotated coordinate system respectively aligned with the lateral direction and movement extent. The origin of this coordinate system was the center of the virtually displayed targets.

### Quantifying Exploratory Behaviour with Lag-1 Autocorrelation

Following the approach of van Beers and colleagues,^8^ we quantified the level of exploration along the solution manifold by calculating lag-1 autocorrelations on trial-by-trial final hand position. Here, larger correlation values suggest greater exploratory behaviour. Recently, we postulated that reinforcement feedback that alters reach aim may give rise to lag-1 autocorrelations,^18^ but this idea had not yet empirically tested. For each experiment, we performed autocorrelation analysis separately along the major and minor axes of the rectangular targets. In the **Results**, we focus on lag-1 autocorrelations along the major axis of the targets to compare between task-redundant and task-relevant dimensions.

### Reinforcement-Based Models of Final Hand Position

The majority of sensorimotor reaching and final hand position models consider no movement variability^67,68^ or a single source of movement variability.^15,69^ Only a handful of error-based^31,8^ and reinforcement-based^18,39,32^ models of final hand position consider multiple sources of movement variability. To our knowledge, three different sources of movement variability have been reported in the literature: i) motor movement variability,^25,26,27^ ii) planned movement variability,^8,34^ and iii) exploratory movement variability.^22^ Note that motor movement variability has also been referred to as “motor noise”^15^ or “execution variability”.^27^

#### Model 1

Reinforcement-based models of final hand position have used various assumptions regarding how the sensorimotor system updates its reach aim following feedback and how much knowledge the sensorimotor system has of a particular variability source. Here we have developed a general set of equations (Model 1), using a minimal set of assumptions that consolidates past models of reinforcement-based models of final hand position. We used this general model to make *a priori* predictions of 2D endpoint reach behaviour and resulting exploratory behaviour. Here we modeled 2-dimensional endpoint reach position (*X_t_*) and intended reach aim 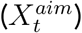 as

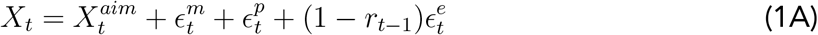

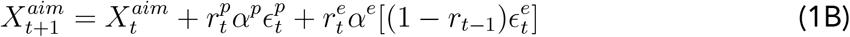

Final reach position on the current trial (*X_t_*) is equal to the intended reach aim 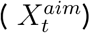 with additive Gaussian noise 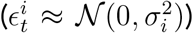. Superscripts represent the source of the variability: motor movement variability (*m*), planned movement variability (*p*) and exploratory movement variability (*e*). In the equations, variability terms 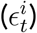 are covariance matrices with an y (movement extent) and x (lateral position) component. Note that exploratory movement variability is added only if the previous trial was unsuccessful^18,30,39,32^ 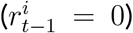. If the current trial is successful 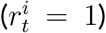, reach aim is updated proportionally (*α_p_,α_e_*) to planned and exploratory movement variability.

We used Model 1 to make *a priori* predictions of individual level and group level behaviour. Individual behaviour is predicted by using movement variability to update its intended reach aim based on positive reinforcement feedback. Updating reach aim towards a recently successful movement results in trial-by-trial random-walk exploration of the solution manifold. Performing lag-1 autocorrelation analysis on the simulated hand positions within each condition allowed us to quantify the level of exploration. Group level predictions are generated by simulating 18 individuals. We then averaged the resulting lag-1 autocorrelations across the simulated individuals.

The general model (Model 1) may not be the simplest model that can explain our experimental data. Further, the general model has a large number of free parameters and may consequently be prone to overfitting. To find the simplest model, we systematically reduced the model’s free parameters and assumptions. Each model derivation proposes a slightly different interpretation of the mechanism regulating exploration of the solution manifold. We used Bayesian Information Criteria and Akaike Information Criteria analyses to test how well each captures the underlying sensorimotor process. We considered 6 additional models. Models 2-5 are derived from our general model (Model 1). Models 6 and 7 consider reinforcement-based models previously proposed in the literature.^18,32^

#### Model 2 (Remove 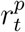)

Our general model (Model 1; **Eq. 1A, 1B**) assumes that updating intended reach aim using planned and explorative movement variability is conditioned on reinforcement feedback. However, it is possible that updates to reach aim using planned movement variability are not conditioned on reinforcement feedback, but are rather always present.^8^ Thus, we considered the following model that always updates its reach aim using planned movement variability, regardless of reinforcement feedback:

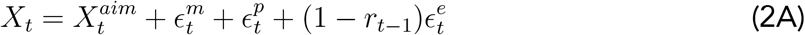

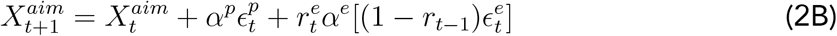

#### Model 3 (Remove 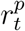, *α^p^*)

The general model (Model 1; **Eq. 1A, 1B**), updates using planned movement variability and exploratory movement variability are considered as separate processes by using separate update terms. This formulation would suggest that two separate neural systems control planned and exploratory movement variability. However, another possibility is that the same neural system governs both these processes. Here we consider the idea that a single process, using a single update term, governs reach aim updates when using planned and exploratory movement variability:

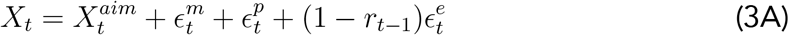

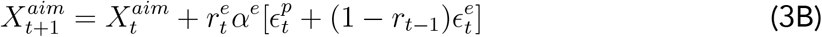

#### Model 4 (Remove 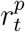, *α^p^, ϵ^p^*)

Another possibility is that planned movement variability is not used to update intended reach aim. That is, only exploratory movement variability is used to update reach aim following successful trials. Here we remove all terms related to planned movement variability. The resulting model captures the idea that a single reinforcement-based process regulates exploration of the solution manifold.

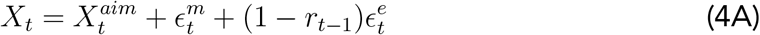

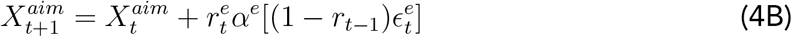

#### Model 5 (Remove 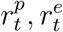, *α^e^, ϵ^e^*)

Previous work from van Beers and colleagues^8^ studied exploration along the task-redundant, which they termed task-irrelevant, solution manifold during an error-based task. They proposed an error-based model that produced random-walk exploration along the task-redundant dimension. Their model did not rely on reinforcement-based mechanisms. Rather, their model generates random-walk behaviour by both allowing planned movement variability to accumulate and tuning their error correction term separately for the task-redundant and task-relevant dimensions. Similarly, we consider a model that accumulates planned movement variability without the use of reinforcement-based mechanisms.

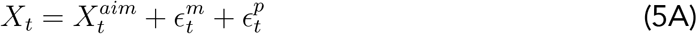

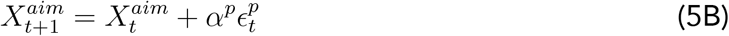

Note this model closely resembles the use-dependency hypothesis.^55,70^ The use-dependency hypothesis suggests that the sensorimotor system will bias the next movement to be similar to the previous movement. Aligning with Model 5, the use-dependency hypothesis also suggests this behavioural change is not conditioned on positive reinforcement. However, it should be noted that Model 5 differs from the original use-dependency model, which does not include planning movement variability.

#### Therrien 2018

We considered a model proposed by Therrien and colleagues (2018) that utilizes complete knowledge of exploratory movement variability to update its intended reach aim after a successful trial. To this end, they used an exploratory movement variability term that was always present but increased in variance following a failure. This formulation is equivalent to having planned movement variability that is always present and using additional exploratory movement variability following failure, which is more consistent with the terminology and notation used in this manuscript. Using this notation, we have rewritten the Therrien 2018 as

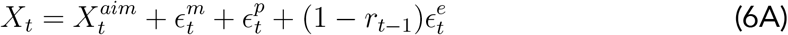

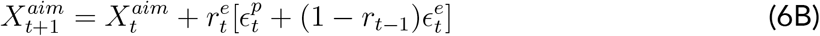

Note that although we have adapted the original notation to align with our terminology and notation, the equation above is mathematically equivalent to the original Therrien 2018 model.

#### Cashaback 2019

We recently proposed a reinforcement-based model of final hand position to understand the role of the underlying reinforcement gradient on the rate of learning.^18^ This model used a portion of motor movement variability and exploratory movement variability to update its intended reach aim after a successful trial. This model can be expressed with the current notation as

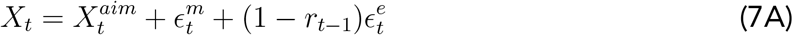

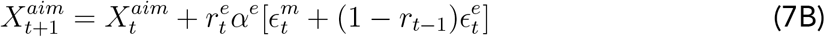

### Model Fitting

Seven different models were considered in total. We used the same fitting procedure for each of these seven models. Model fitting was performed using the Powell algorithm in the Minimize function from the Scipy Python library.

For each model, we simulated 500 participants in each experiment. We then calculated the mean lag-1 autocorrelation for each condition along both the major and minor axes of the rectangular targets. For each condition, we defined the loss function as the absolute difference between the average simulated and average participant lag-1 autocorrelation along the major and minor axes of the target. The optimizer simultaneously minimized the sum of the loss across all three experiments.

The fitting procedure began with a “warm-start” procedure. Here, we minimized model loss using randomized initial parameter guesses. We repeated this process 1,000 times. From these 1,000 randomized initializations, we then selected the set of parameters that resulted in the lowest loss. Using these parameters, we then performed a bootstrapping procedure (100,000 iterations) to find the 95% confidence intervals for each free parameter. Each bootstrap iteration used the same parameters from the warmstart procedure. In each bootstrap iteration, for each condition we resampled with replacement participant lag-1 autocorrelations. The mean lag-1 autocorrelation from this resampled group was used in determining model loss for each bootstrap. Update terms (p, e) were bounded from 0 to 1. The variances used to estimate the sources of movement variability had a lower bound of 0. This bootstrap procedure estimates the posterior distribution of each parameter.

### Best-Fit Model Selection

We found the median from the corresponding posterior distribution for each parameter. Using these median parameter values (Θ), we simulated each condition with each model (Models 1-7). Again, across all three experiments we used a loss function (L) that took the absolute difference of between the average simulated (*ACF^Model^*) and average participant (*ACF^Data^*) lag-1 autocorrelation (*ACF*) along the two target axes (i; major and minor) in each condition (j):

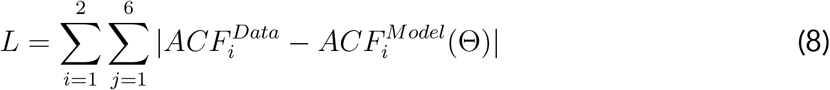

Thus, a model that produces lag-1 autocorrelations resembling the data will have a low final loss. This loss was used in Bayesian Information Criteria and Akaike Information Criteria analyses to select the model that best captured the data with the fewest parameters. Bayesian Information Criteria (**Eq. 9**) weights the number of free parameters (k) by the number of datapoints (n) used to calculate the loss (L). However, Bayesian Information Criteria (BIC) can be biased towards models with fewer parameters. We also considered Akaike Information Criteria (**Eq. 10**) which uniformly weights free parameters (k) used in the model. Akaike Information Criteria (AIC) can be biased towards models with more free parameters. For both BIC and AIC analyses, a lower score indicates a better fit. We considered both analyses when selecting the best-fit model. We defined our best-fit model, as the model that yielded the lowest Bayesian Information Criteria and Akaike Information Criteria scores.

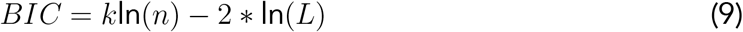

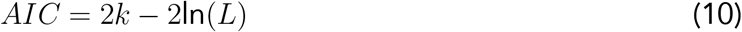

Overall, the model fitting and comparison analysis took approximately 120 hours to perform on an 8 core Intel i7-107000 processor. Parameter posterior probability distributions for the best-fit model can be found in **Supplementary C**.

### Average Reach Position

In **Experiment 2**, for each participant we compared their average final hand position during the hidden long rectangular reward zone condition to their reported reach aim. For each participant we estimated the 95% confidence interval of their average final hand position with a bootstrapping procedure. We defined the participant’s average reach to be outside the target if the entire 95% confidence interval is outside the virtually displayed target.

### Shuffled Lag-1 Autocorrelation

We hypothesized that the sensorimotor system uses a reinforcement-based mechanism to explore the task-redundant solution manifold. Theoretically, reinforcement-based mechanisms would be sensitive to our experimental manipulations and exhibit different levels of exploration depending on condition. Specifically, a reinforcement-based mechanism would result in a different sequential ordering of final hand positions and different levels of exploration (lag-1 autocorrelations) for different experimental conditions. However, we wanted to control for the possibility that the observed lag-1 autocorrelations were a result of some unknown stochastic processes (e.g., pink noise; power signal density is inversely proportional to signal frequency) rather than the sequential ordering of final hand positions based on reinforcement feedback. Note that a purely gaussian process (i.e., white noise: equal power signal density of all signal frequencies) with no update toward successful actions would result in a zero lag-1 autocorrelation and would be unable to explain our results. We performed a shuffling analysis^12,41^ to control for unknown stochastic processes causing the observed lag-1 autocorrelations. For the shuffling analysis, we defined the null hypothesis to be that a participant’s observed lag-1 autocorrelation was an artefact of unknown stochastic processes.

To estimate the null hypothesis, we created a shuffled array of final hand positions by randomly sampling without replacement for each participant per condition. Thus, the resulting shuffled array had the same mean and standard deviation of the original dataset, but a different sequential ordering of final hand positions. We then calculated the lag-1 autocorrelation of the shuffled array. For each participant per condition, this process was repeated 1,000,000 times. We rejected the null hypothesis if less than 5% of the shuffled arrays had an autocorrelation as large or larger than the participant’s experimentally observed autocorrelation. Rejecting the null would support the idea that the observed lag-1 autocorrelations in each condition were due to reinforcement feedback and not an unknown stochastic process.

### Movement Variability, Trial-by-Trial Difference

We^18^ and others^62,30,39^ have found that binary reinforcement feedback modulates movement variability. We calculated movement variability along the dimensions aligned with the major and minor axes of the rectangle targets during the baseline and experimental conditions. Movement variability was calculated as the standard deviation of the trial-by-trial change (ΔX^*i*^) in final hand positions. Baseline movement variability was calculated during the last 10 trials of the baseline condition. Experimental condition movement variability was calculated independently for reinforced and unreinforced reaches.

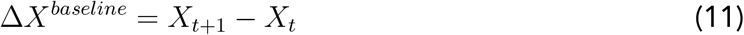

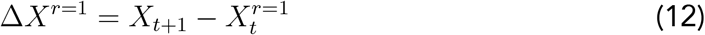

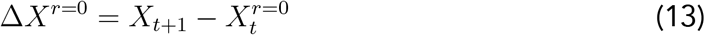

Here, *X* represents a participant’s final reach position, *t* is the trial number, and r represents if the reach was reinforced on the current trial (1 = reinforced, 0 = unreinforced).

### Movement Variability, Distribution of Final Hand Positions (IQR)

Past studies have quantified exploration by comparing the relative variability between redundant and relevant solution manifolds^4,11,6,7^ (i.e., ‘uncontrolled manifold’ and ‘orthogonal dimension’). We quantified the magnitude of movement variability of final hand position by calculating the interquartile range (IQR). IQR is known to be a robust measure of variability^71^ because it is not heavily influenced by outliers, unlike other measures of dispersion such as variance or standard deviation. We calculated IQR as the difference between the 25th and 75 percentiles. For each experiment, we took the IQR ratio between conditions to describe the relative movement variability in final hand position. A value of one represents equal movement variability of final hand position between conditions. In **Experiment 1** and **Experiment 2**, we looked at the ratio between the task-redundant and task-relevant conditions. In **Experiment 3**, we took the ratio between the 80% and 20% probability of reinforcement conditions. Statistical comparisons were made between the mean variability in each experiment and an IQR equal to one. We repeated this analysis using simulations from our best-fit model.

### Reaction Time

Recent literature has examined the effects of reinforcement (reward) feedback and task success on movement vigor.^47,48,50,51^ These studies use reaction times and movement times as proxies of movement vigor. Specifically, a faster reaction time or movement time corresponds with an increase in movement vigor. Reaction time (ms) was calculated as the time between the start position turning yellow (signaling the start of the reach) and participants leaving the start position. We compared participant reaction times following reinforced and unreinforced reaches within each condition.

We did not examine movement times because participants were free to reach to any position along the target. Participants would likely take longer to reach further distances. This would potentially confound our ability to discern whether changes in movement time were due to vigor or reach distance.

### Statistical Analysis

Non-parametric bootstrap hypothesis tests (1,000,000 iterations) were used for followup mean comparisons. We used directional tests when testing theory-driven predictions, and nondirectional tests otherwise. Spearman Rank correlation was used for all correlation analyses. Fisher’s Exact test was used to test for significant differences between reported and actual reach aim. We computed common language effect sizes 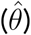 for all mean comparisons. Statistical tests were considered significant at p < 0.05.

## SUPPLEMENTARY

### Supplementary A: Lag-1 Autocorrelations Along the Minor Axis

**Supplementary Figure S1:**
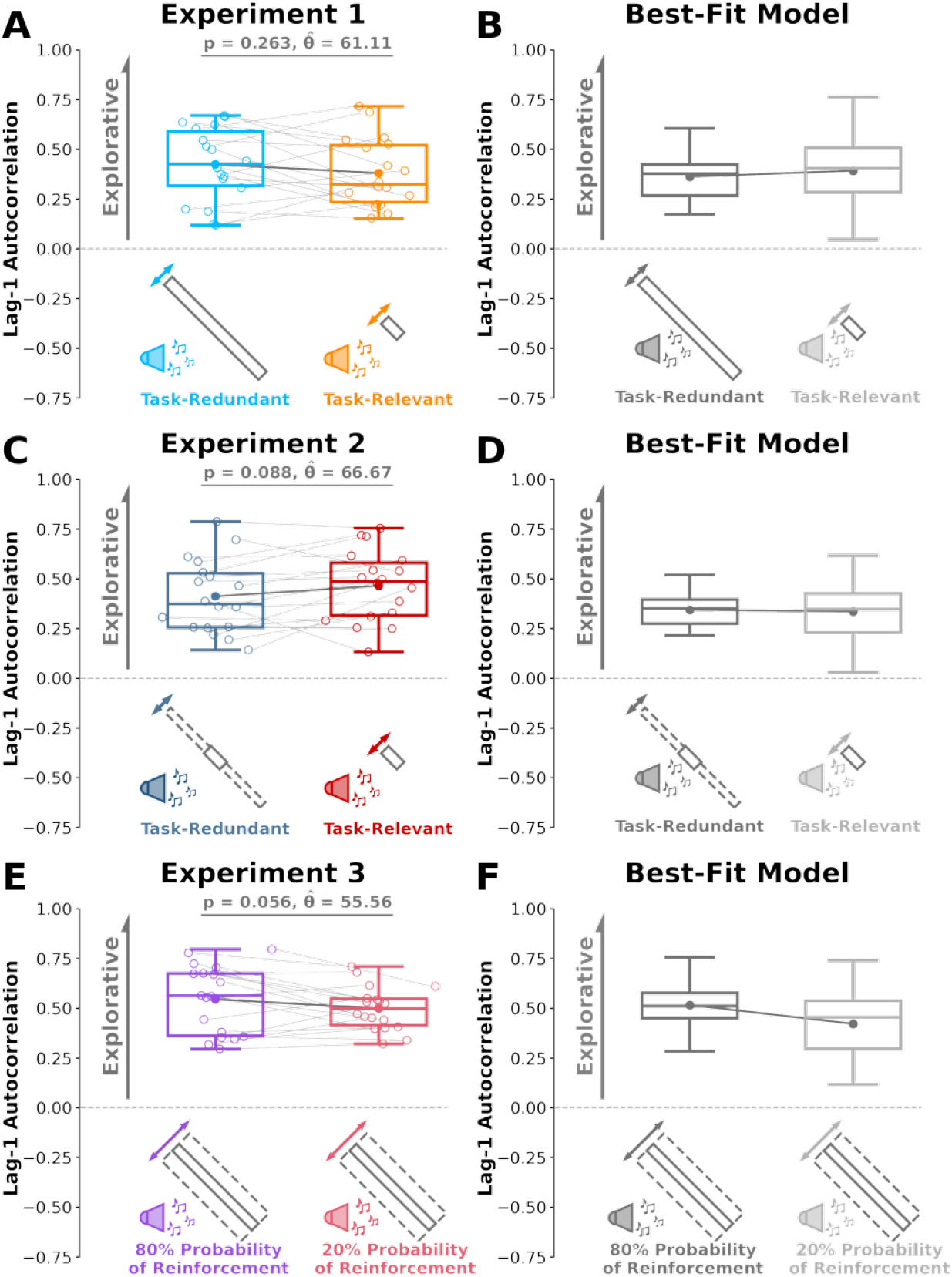
Apparatus and Experiment Design. **A)** We calculated lag-1 autocorrelation (y-axis) along the minor axis of the rectangular targets for each condition (x-axis). Solid circles and connecting lines represent mean lag-1 autocorrelation for each condition. Box plots represent the 25th, 50th, and 75th percentiles. We did not find significant differences in lag-1 autocorrelation along the minor target axis between conditions across **A) Experiment 1**, **C) Experiment 2**, or **E) Experiment 3**. Corresponding lag-1 autocorrelations from the best-fit model for **B) Experiment 1**, **D) Experiment 2**, or **F) Experiment 3**. The best fit model did well to capture the absolute levels of lag-1 autocorrelations. We did not expect differences between conditions in **Experiment 1** and **2**. Our model did predict differences between the 80% and 20% conditions along the minor axis in **Experiment 3**, but participants did not display significant paired differences despite trending in the expected direction (p = 0.056).

### Supplementary B: Shuffling Analyses to Control for Potential Spurious Stochastic Processes

We hypothesized that the sensorimotor system uses a reinforcement-based mechanism to explore the task-redundant solution manifold. Specifically, we tested the idea that a reinforcement-based mechanism would result in a sequential ordering of trial-by-trial final hand positions that resulted in different levels of exploration (lag-1 autocorrelations) according to an experimental condition. However, we wanted to control for the possibility that the observed lag-1 autocorrelations were a result of unknown stochastic processes rather than the sequential ordering of final hand positions based on reinforcement feedback.

To test whether unknown stochastic processes caused the observed lag-1 autocorrelations, we performed a shuffling analysis^12,41^ (see **Methods**). For each participant in each experiment, we shuffled their trial-by-trial final hand positions within each condition. This shuffled the ordering of final hand positions while keeping the statistical moments (i.e., mean, variance, etc.) constant. We then calculated the lag-1 autocorrelation of the newly shuffled hand positions. This process was repeated for 1,000,000 iterations. Here the null hypothesis is that the observed lag-1 autocorrelations were the result of an unknown stochastic process. We rejected the null hypothesis if less than 5% of these shuffling iterations resulted in a lag-1 autocorrelation larger than the originally observed lag-1 autocorrelation. We concluded that the original lag-1 autocorrelation for each participant in each condition was not an artefact of unknown stochastic processes (p < 0.05 for all participants). This analysis further supports the idea that the sequential trial-by-trial ordering of final hand position was caused by the experimental manipulations.

**Supplementary Figure S2:**
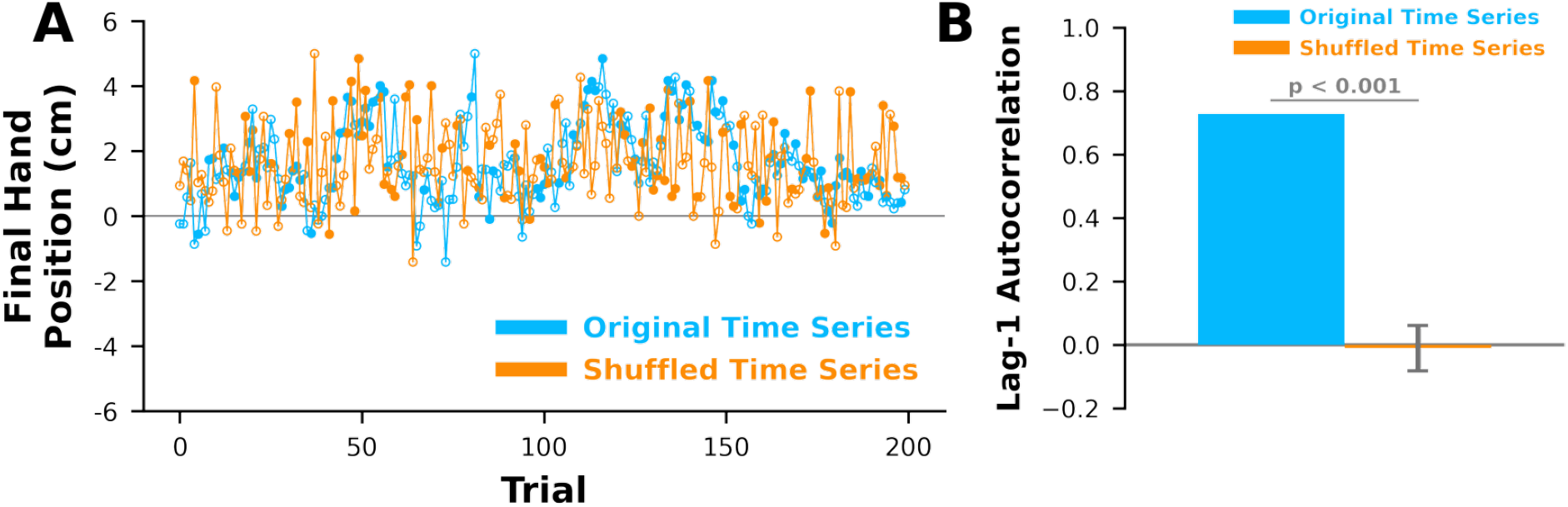
Shuffled Final Hand Positions. Here we show the shuffling analysis for a representative participant. **A)** Final hand positions (y-axis) over the trials (x-axis) for the original time series (light blue) and shuffled version of the exact same time series (light orange). We shuffled the original time series such that the statistical moments (i.e. mean, variance, etc.) were held constant by rearranging the trial-by-trial ordering of final hand positions. **B)** We calculated the lag-1 autocorrelation (y-axis) of the shuffled time series (light orange) and compared it to the original time series (light blue). Error bars represent standard deviation. This process was repeated 1,000,000 times. We concluded that a spurious stochastic process could not explain the original time series if less than 5% of the shuffled time series were as large or larger than the original time series (p < 0.05 for all participants).

### Supplementary C: Best-Fit Model Parameters

**Supplementary Figure S3:**
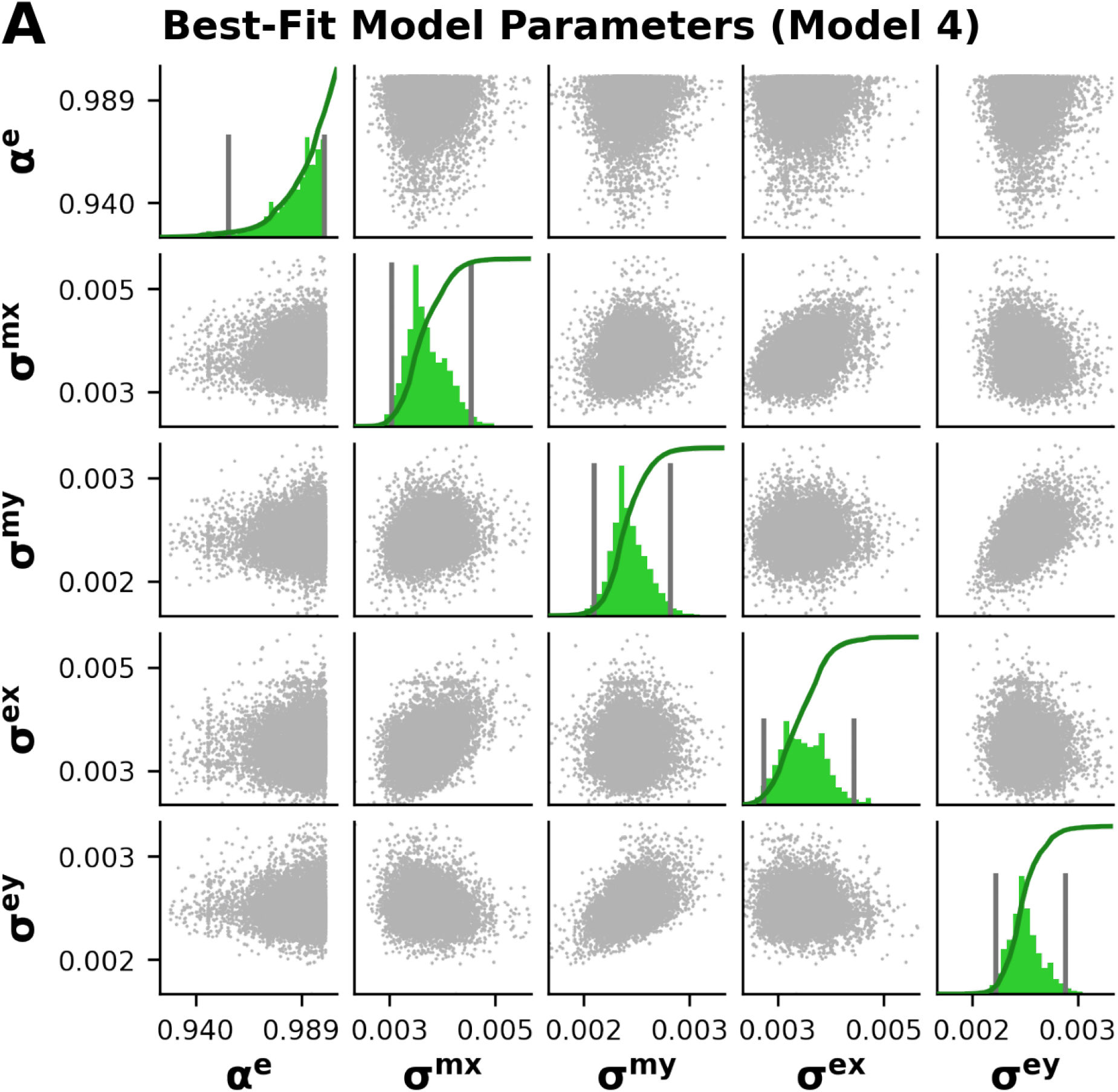
Best-fit Model Parameter Distribution. **A)** Model parameters for the best-fit model (Eq. 4, Model 4). Bootstrapped (10,000 samples) marginal probability distributions of each parameter value are shown along the diagonal (green). 95% confidence intervals are indicated with dark grey lines. Thick dark green lines correspond to the cumulative distribution. We used the median value for *α^e^*(0.99), *σ^mx^* (3.93 mm), *σ^my^*(2.29 mm), *σ^ex^* (3.40 mm), and *σ^ey^*(2.18 mm) to run simulations with the best-fit model. Off-diagonal plots show joint probability distributions for each pair of parameters.

### Supplementary D: Movement Variability

**Supplementary Figure S4:**
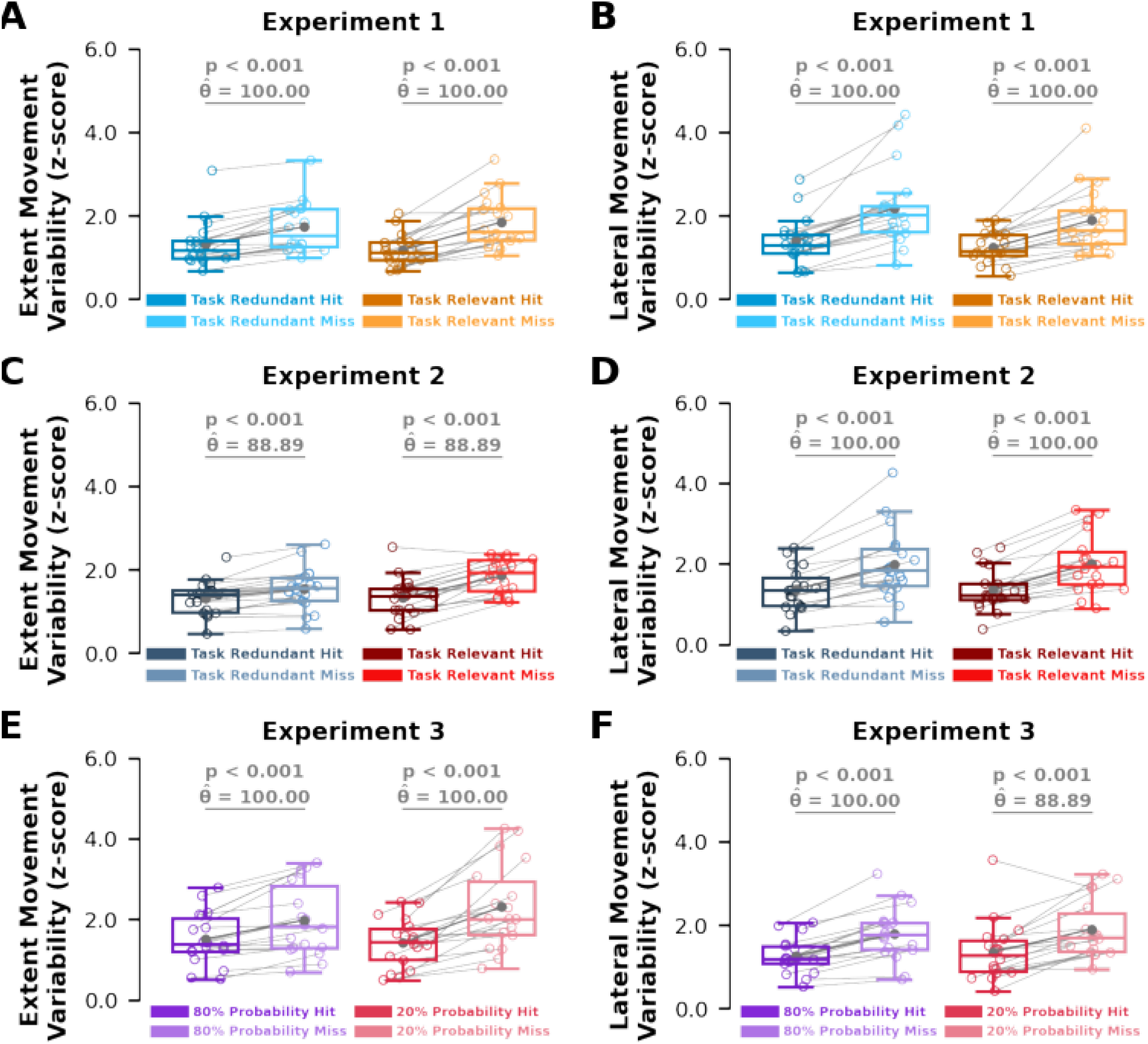
Movement Variability. We calculated movement variability in each condition separately following successful trials (Hit, dark colours) and unsuccessful trials (Miss, light colours). Here, we defined movement variability as the standard deviation of the trial-by-trial change in reach position. Reach position coordinates were projected onto a coordinate system aligning with the movement extent (left column) and the lateral direction(right column). We normalized this variability to individual baseline movement variability to calculate a z-score (y-axis) for each participant. Solid circles and connecting lines represent mean movement variability for each condition. Box plots represent the 25th, 50th, and 75th percentiles. We found participants displayed significantly increased movement variability following an unreinforced trial (Miss, light color) compared to reinforced trials (Hit, dark color) across **A,B) Experiment 1**, **C,D) Experiment 2**, and **E,F) Experiment 3**.

### Supplementary E: Movement Vigor

Recent literature has examined the effects of reinforcement (reward) feedback and task success on movement vigor.^47,48,49,50,51^ Specifically, they showed that faster reaction times correspond to higher movement vigor. We compared reaction times following reinforced trials and unreinforced trials within each condition across all three experiments (**Fig. S4**). Reaction time was defined as the time between the start target turning yellow (signifying the beginning of the trial) and the participant leaving the start position. We found a significantly greater reaction time following reinforced trials compared to unreinforced trials in the task-redundant condition from **Experiment 1** (p = 0.039, = 72.22). However, we did not find this relationship for the remaining five conditions (p >= 0.147, *θ* <= 67.11 for all other comparisons). Thus, we did not find consistent evidence across all three experiments that suggests reward leads to decreased reaction times during exploratory behaviour.

**Supplementary Figure S5:**
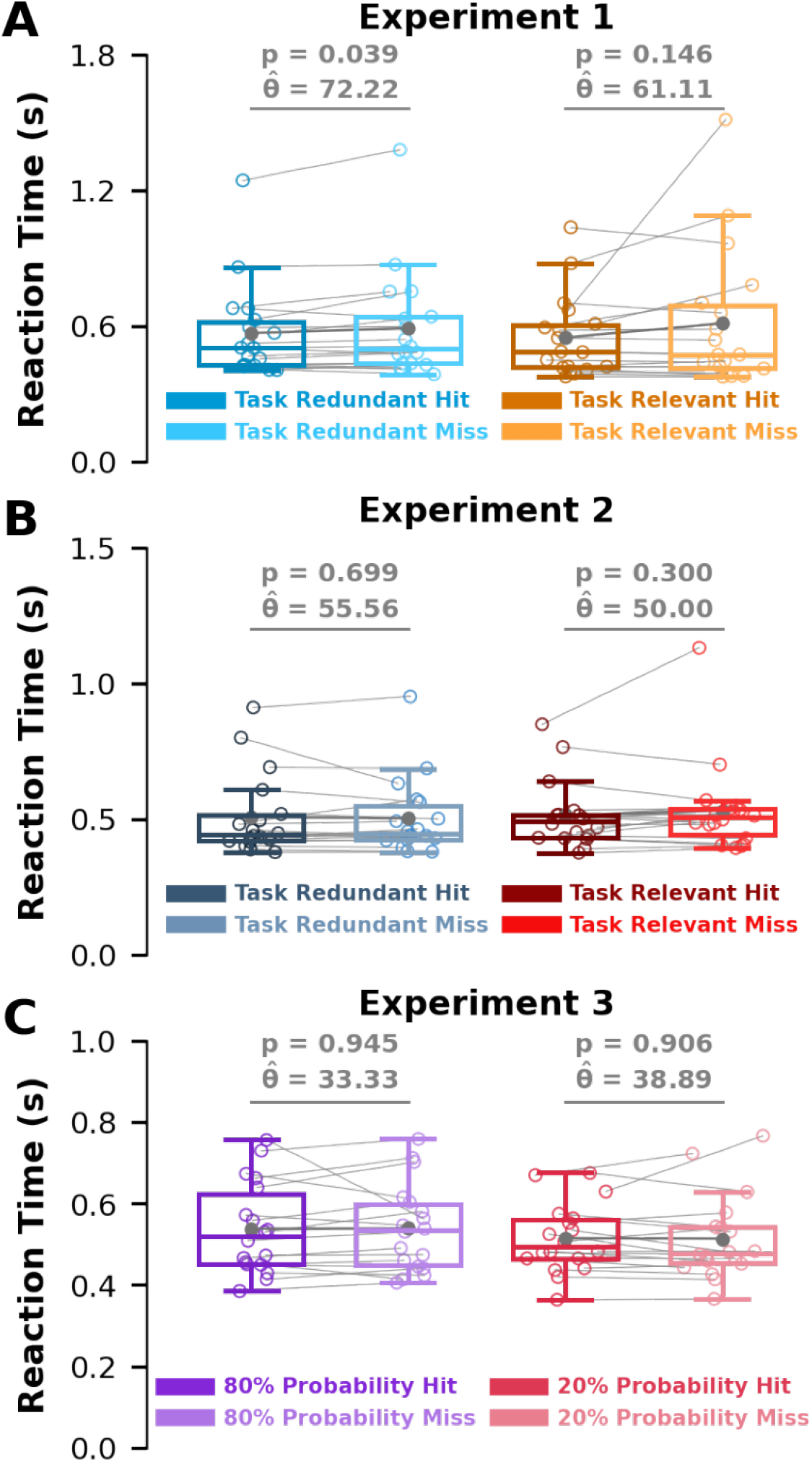
Reaction Time. We calculated participant reaction time (y-axis) as the time for a participant to leave the start position after the start position changed colour. **A)** We found that participants had a significantly faster reaction time (p = 0.039) after a successful trial (Hit) compared to an unsuccessful trial (Miss) in the task-redundant condition (light blue) of **Experiment 1**. However, this finding was not found in the task-relevant condition (light orange). We found no significant difference in reaction times between successful trials and unsuccessful trials in **B) Experiment 2** or **C) Experiment 3**.

## Notes

### Competing Interest Statement

The authors have declared no competing interest.

### Summary of Updates

Minor updates have been made to terminology for better clarity and accuracy.

